# Airway tissue stem cells reutilize the embryonic proliferation regulator, Tgfß-Id2 axis, for tissue regeneration

**DOI:** 10.1101/2020.11.23.394908

**Authors:** Hirofumi Kiyokawa, Akira Yamaoka, Chisa Matsuoka, Tomoko Tokuhara, Takaya Abe, Mitsuru Morimoto

## Abstract

During development, quiescent basal stem cells are derived from proliferative primordial progenitors through the cell cycle slowdown. In contrast, quiescent basal cells contribute to tissue repair during adult tissue regeneration by shifting from slow-cycling to proliferating and subsequently back to slow-cycling. Although sustained basal cell proliferation results in tumorigenesis, the molecular mechanisms regulating these transitions remain unknown. Using temporal single-cell transcriptomics of developing murine airway progenitors and *in vivo* genetic validation experiments, we found that Tgfß signaling slowed down cell cycle by inhibiting *Id2* expression in airway progenitors and contributed to the specification of slow-cycling basal cell population during development. In adult tissue regeneration, reduced Tgfß signaling restored *Id2* expression and initiated epithelial regeneration. *Id2* overexpression and *Tgfbr2* knockout enhanced epithelial proliferation; however, persistent *Id2* expression in basal cells drove hyperplasia at a rate that resembled a precancerous state. Together, the Tgfß-Id2 axis commonly regulates the proliferation transitions in airway basal cells during development and regeneration, and its fine-tuning is critical for normal regeneration while avoiding basal cell hyperplasia.

## Introduction

Tissue stem cells contribute to homeostasis through the strict regulation of cell proliferation (Leach and Morrisey, 2018; Yanger and Stanger, 2011). Under normal conditions, most of the tissue stem cells are maintained in the quiescent or slow-cycling state by suppression of cell cycle progression (Furutachi, et al., 2015; Desai, et al., 2014; Cheung and Rando, 2013). However, following injury, they undergo self-renewal by re-entering the cell cycle for tissue regeneration (Cheung and Rando, 2013; Yanger and Stanger, 2011). The precise regulation of the transitions of the stem cell population between the quiescent/slow-cycling and active cell cycle is essential for tissue homeostasis; the dysregulation of this transition is related to pathological disorders, such as cancer (Feitelson, et al., 2015; Lapouge, et al., 2011; Barker, et al., 2009). However, the molecular mechanism regulating the stem cells’ transitions to the proliferation mode remains unknown. Thus, understanding such molecular mechanism is important for stem cell biology and human health.

Quiescent tissue stem cells are selected and segregated from the primordial progenitors during tissue development. It has been reported that the tissue stem cells of hair follicles and the brain enter the quiescent/slow-cycling state as they differentiate from the primordial progenitors during organogenesis (Furutachi, et al., 2015; Shyer, et al., 2015; Gancz, et al., 2011; Nowak, et al., 2008; Mikkola and Orkin, 2006). These studies have demonstrated the importance of cell cycle slowdown for the specific quiescent tissue stem population at the early stage of tissue development. For example, in the development hair follicle stem cells, slow-cycling cells expressing stem cell markers appear in the primordial surface ectoderm. This population differentiates into mature quiescent hair follicle stem cells at the later stage. The ablation of this slow-cycling population compromised epidermal would repair due to the decreased number of tissue stem cells (Nowak, et al., 2008). Thus, cell cycle slowdown within primordial progenitors must be critical for the specification of tissue stem cell population during development. However, the molecular mechanism inducing the cell cycle attenuation in the progenitor population remains largely unknown.

Adult airway epithelium shows a low cellular turnover rate, usually more than 4 months, in rodents (Rock, et al., 2009; Blenkinsopp, 1967). It possesses a substantial ability to regenerate damaged cells in response to severe injuries caused by a viral infection or chemical toxicity (Hogan, et al., 2014; Kotton and Morrisey, 2014; Rock, et al., 2011). Airways display pseudostratified epithelium, which comprises four major cell types, including ciliated, club, neuroendocrine (NE), and basal cells (Herriges and Morrisey, 2014; Morrisey and Hogan, 2010). Airway basal cells are well-characterized epithelial tissue stem cells (Pardo-Saganta, et al., 2015; Rock, et al., 2009; Hong, et al., 2004) whose capacity for self-renewal is strictly regulated so that they remain as slow-cycling basal cells for more than 16 weeks (Rock, et al., 2009). The dysregulation of airway basal cells can result in fatal diseases, including squamous cell carcinoma (Lapouge, et al., 2011). Sulfur dioxide (SO_2_) inhalation-induced tissue injury is known as an experimental model for airway epithelial injury-regeneration; in this model, slow-cycling basal cells can be activated to proliferate and produce transit-amplifying cells within 48 hours. After these proliferating transit-amplifying cells repair the epithelial damage, they subsequently start differentiating into luminal cells to stop the cell cycle progression and hyperplasia (Pardo-Saganta, et al., 2015; Rock, et al., 2011).

During airway development, the basal cells are selected and segregated from primordial airway progenitors, which show high proliferation profile and arise from the ventral foregut (Herriges and Morrisey, 2014; Morrisey and Hogan, 2010). During lung development, primordial airway progenitors commit to the cell lineage, slowing down the cell cycle and acquiring mature cells’ canonical markers (Herriges and Morrisey, 2014). We had previously reported the crucial function of Notch signaling in alternative cell fate selection in club, ciliated, and NE cells (Kiyokawa and Morimoto, 2020; Arner, et al., 2015; Consortium, et al., 2014; Morimoto, et al., 2012; Morimoto, et al., 2010). However, the specific mechanism of slow-cycling basal progenitors during development remains unclear.

In the present study, to unveil the molecular mechanism that establishes slow-cycling basal cells, we delineated a comprehensive developmental roadmap of mouse airway epithelial cells using time series, single-cell transcriptome analyses. This approach and *in vivo* lineage tracing experiments defined the trajectory of basal cell differentiation and identified novel *Krt17^+^* basal progenitors. We also found that genes encoding inhibitors of DNA-binding/differentiation (Id) proteins promote epithelial proliferation. Tgfβ signaling reduces *Id2* expression to slow down the progenitors’ cell cycle around E14.5, inducing the *Krt17^+^* basal progenitors and *Scgb3a2^+^* luminal progenitors.

In addition, we demonstrated that the Id2 dosage regulates the proliferation of mature basal cells. At the perinatal and adult stages, *Id2* expression is further restricted and maintained at low levels only in mature basal cells to ensure their proliferation potential. The SO_2_-exposure mediated injury-regeneration model using adult mice showed that an increased *Id2* expression by Tgf*β* suppression initiated the regeneration of the injured epithelia via promoting basal cells’ re-entry into the cell cycle. Furthermore, the artificial enhancement of *Id2* expression boosted basal cell proliferation and resulted in basal cell hyperplasia, which resembles a precancerous state. These results imply that the fine-tuning of *Id2* expression is critical for normal tissue regeneration while avoiding tumorigenesis.

In summary, we demonstrate that the Tgfß-Id2 axis is a shared, critical regulator of the transition between the active proliferation and slow-cycling mode in airway stem cells during development and adult tissue regeneration.

## Results

### Time series single-cell RNA sequencing (scRNA-seq) analyses to delineate a developmental roadmap of airway epithelial cells, including basal cells

We aimed to identify a high-fidelity marker of early basal progenitors to elucidate the developmental process of mature basal cells; p63 is a well-known marker for basal cells but is not restricted to basal progenitors at early development (Yang, et al., 2018). We used a droplet-based scRNA-seq with approximately 3500 epithelial cells at six time points from E12.5 to E18.5 (Figure 1A). This approach allowed us to delineate a comprehensive lineage map of airway epithelial cells derived from respiratory endoderm during embryogenesis. We visualized the distinct populations with cluster analysis using the t-SNE algorithm. Uniform progenitors at E12.5 and E13.5 changed transcriptome profiles at E14.5 and acquired cell type-specific gene signatures by E16.5 (Figures 1B and S1B, also see Figure 3I). During E12.5 to E14.5, proliferative markers are frequently detected and are the major determinants for cell clustering analysis (Figure 1B and see Figure 3A). Epithelial progenitors eventually differentiated into three major populations, basal, club, and ciliated cells, and one minor population, NE cells (Figure 1B). This transcriptome clustering and the pseudo-time trajectory analysis using Monocle (Figure 1C) demonstrated that the lineage-specific transcriptional patterns were determined between E14.5 and E16.5. *Foxj1^+^* ciliated cells first appeared at approximately E14.5, which was suggested because some of the E14.5 progenitors were classified in the ciliated cell population (Figure 1B). Other progenitors differentiated into *Krt5^+^* basal cells or *Scgb1a1^+^* club cells, beginning segregation at E15.5 and becoming more apparent after E16.5 (Figure 1C). The detection of gene signatures of NE cells, such as *Ascl1* and *Cgrp*, at E13.5 (Figures 1D and S1A) showed that the differentiation of NE cells occurred the earliest among the four cell populations; this finding was consistent with a previous study (Linnoila, 2006).

**Figure 1.**
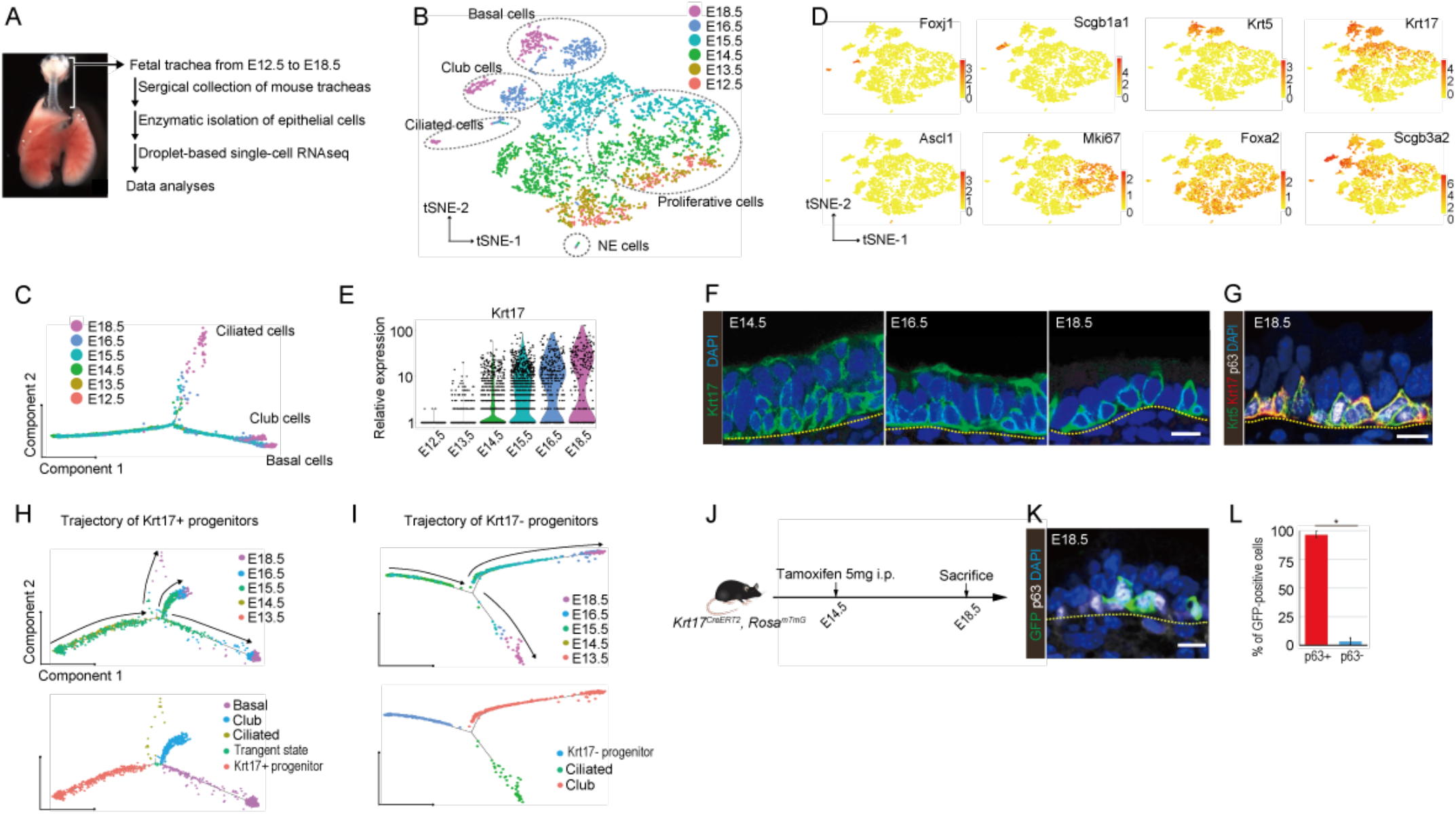
Time-series scRNA-seq analyses delineate a developmental roadmap of airway epithelial cells, including basal cells. (A) Schematic representation of the time-series scRNA-seq analyses of developing tracheal epithelial cells. (B) The t-SNE plot of single cells displayed thirteen distinct clusters, including four distinct mature cell types. (C) Pseudotime analysis with the Monocle package illustrated the developmental trajectories towards three mature cell lineages. (D) Canonical marker expression in the t-SNE map. (E) The temporal expression of *Krt17* in the scRNA-seq data. (F-G) Immunostaining for Krt17 in the developing trachea confirmed the limited expression pattern in basal cells. *In silico* trajectory analyses of the *Krt17*-expressing (H) and nonexpressing progenitors (I) using Monocle. (J) Lineage tracing experiment for the Krt17^+^ progenitors at E14.5 using the *Krt17^CreERT2^ Rosa26^mTmG/+^* mice. (K) Immunostaining for GFP and p63 revealed that most Krt17^+^ progenitors at E14.5 contributed to p63^+^ cells at E18.5 (L) (mean±SD, n=4). * p< 0.05; Student’s t test. Scale bars, 5 μm.

### Binary cell fate decision between *Krt17^+^* basal or *Scgb3a2^+^* luminal intermediate progenitors in primordial progenitors

We sought a novel maker for basal progenitor more committed to the basal lineage than p63-expressing cells, and Krt17 was selected as a candidate (Supplementary Table 1). scRNA-seq data and immunostaining revealed that Krt17 mRNA and protein appear at E13.5 and E14.5, respectively (Figures 1D-F). Krt17-expressing cells become committed to Krt5^+^ basal cells by E18.5 (Figure 1G). We used computational trajectory analyses (Figures 1H and 1I) and indicated that Krt17^+^ progenitors contribute cells to all three major epithelial cells, including basal cells, but Krt17^-^ progenitors do not become basal cells. Therefore, Krt17 expression is a novel maker for intermediate population in the basal cell lineage.

To validate Krt17 as a novel maker for intermediate population in the basal cell lineage, we conducted in vivo lineage tracing experiment with *Krt17^CreERT2^ Rosa26^mTmG/+^* mice (Doucet, et al., 2013). A total of 96.8% ± 2.9% (mean ± SD) of *Krt17^+^* progenitors at E14.5 became mature basal cells at E18.5 (Figures 1J-L), indicating that almost all the *Krt17^+^* cells at E14.5 have already committed toward basal cells. Thus, the expression of *Krt17* is an important step for commitment to mature basal cells after the expression of *p63* (Figure 2G).

**Figure 2.**
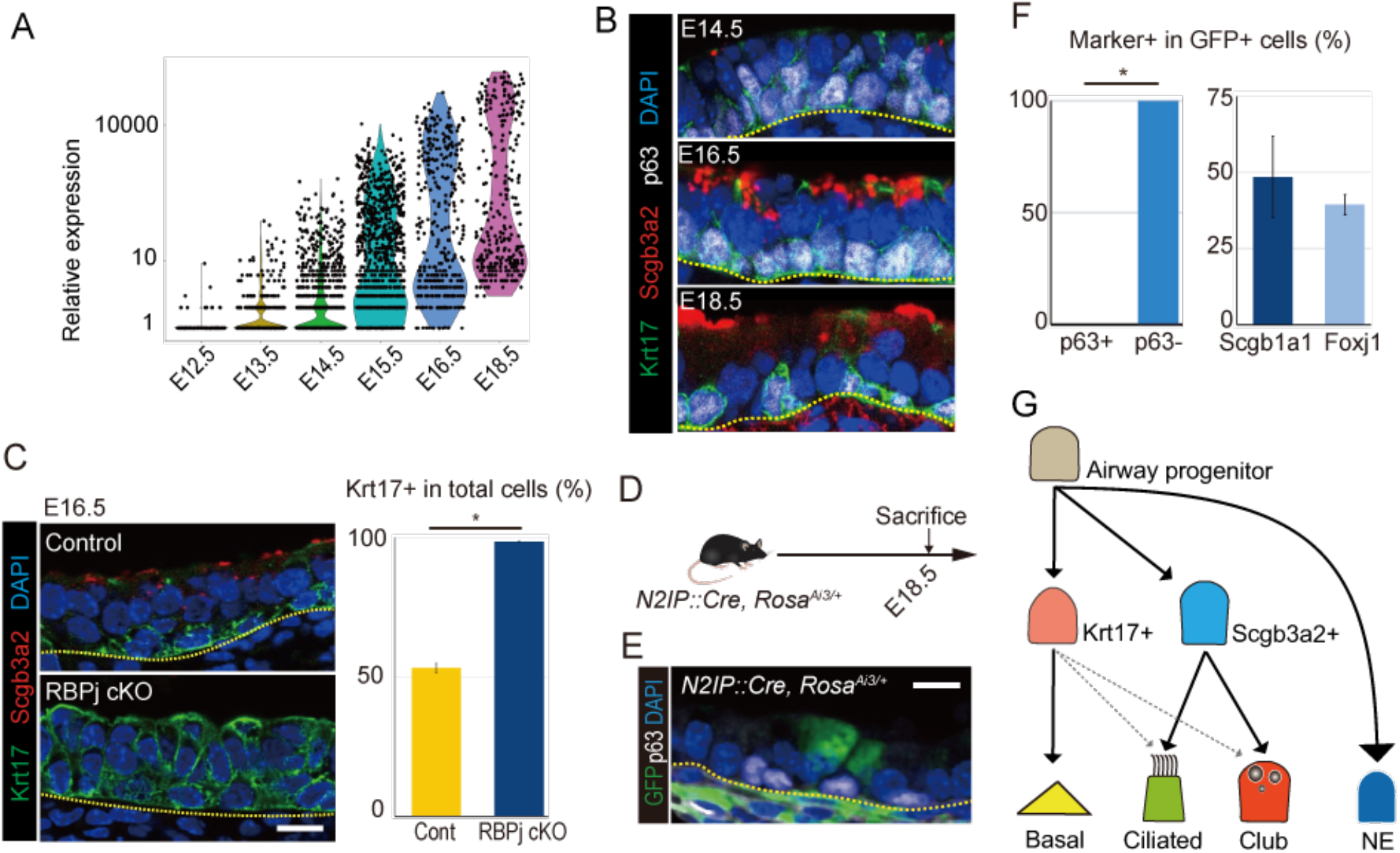
Binary cell fate decision between *Krt17^+^* basal or *Scgb3a2^+^* luminal intermediate progenitors in primordial progenitors. (A) The temporal expression of *Scgb3a2* over time in the scRNA-seq dataset. (B) Actual expression patterns of Scgb3a2 and Krt17 detected by immunostaining. (C) Phenotypic analyses of the RBPj KO mice tracheas at E16.5 by immunostaining for Krt17 and Scgb3a2 revealed the expansion of Krt17^+^ progenitors at the expense of Scgb3a2^+^ progenitors. (D) The lineage tracing experiment for the progenitors that experienced Notch2 activation during development was performed with *N2IP::Cre, Rosa^Ai3/+^* mice. (E,F) Quantitative immunostaining assessment with anti-GFP antibody demonstrated their exclusive contribution to p63^-^ luminal cells at E18.5. (G) Schematic summary of the roadmap for developing airway epithelial cells showing two types of progenitors, Krt17^+^ and Scgb3a2^+^ (Krt17^-^) progenitors. * p< 0.05; Student’s t test. Mean±SD, n=4–6 (C, F). Scale bars: 5 μm.

We further identified Scgb3a2 as a marker gene for Krt17^-^ progenitors by reanalyzing the scRNA-seq data (Figures S2A-C). Scgb3a2 appears from E14.5 on in a pattern mutually exclusive with Krt17 (Figures 2B and S2B), which suggests that equivalent airway progenitors make binary cell fate decision around E14.5 to acquire either Krt17^+^ or Scgb3a2^+^ (Krt17^-^) status that do or do not differentiate into basal cells, respectively. Next, we searched for a cellular signaling pathway that played a critical role in the binary cell fate decision between *Krt17^+^* and *Scgb3a2^+^* progenitors and focused on Notch signaling because Scgb3a2 occurs downstream of it (Guha, et al., 2012). To confirm the role of Notch signaling in the cell fate decision, we genetically ablated Rbpj, a cofactor of Notch signaling, in endodermal epithelium by generating *Shh^Cre^, Rbpj^flox/flox^* mice (RBPj cKO). In this mutant, the complete loss of Scgb3a2 is accompanied by the expression of Krt17 at E16.5 in almost all epithelial cells (Figure 2C). Thus, Notch signaling regulates this binary cell fate decision by inhibiting basal cell specification.

We further asked whether Notch activated cells during development do give rise to luminal cells only, never to basal cells. Given that Notch2 is a major receptor in airway progenitors(Morimoto, et al., 2012), we employed *N2IP::Cre, Rosa^Ai3/+^* mice in which progeny of cells with Notch2 activation expresses a yellow fluorescent protein (EYFP)(Liu, et al., 2013). As expected, EYFP-expressing cells contributed equally to club and ciliated cells but never to basal cells (Figures 2D-F), confirming that Notch-mediated lateral inhibition segregates primordial airway progenitors into Krt17^+^ and Scgb3a2^+^ cells.

Collectively, our time series scRNA-seq analyses show that the gene signature of NE cells is first established at approximately E13.5. The remaining progenitors begin to produce two intermediate progenitors, *Krt17^+^* and *Scgb3a2^+^*, around E14.5 by Notch-mediated lateral inhibition. Basal cell fate is restricted to *Krt17^+^* cells at E14.5, and a mature gene signature such as Krt5 is established at approximately E16.5 (Figure 2G).

### Cell cycle slowdown induces basal cell specification concomitant with downregulation of *Id* gene expression

To elucidate the temporal regulation of epithelial progenitors’ progression into slow-cycling state, we analyzed the scRNA-seq dataset with the Seurat package on the basis of enrichment of cell cycle marker expression. A substantial decrease in numbers of proliferating cells in S and G2-M phases was observed from E14.5 (Figures 3A and 3B), consistent with *in vivo* cell cycle analysis using Fucci mice (*Shh-Cre, Rosa^H2B-EGFP/FucciG1^*). Numbers of cells in G0 phase (quiescent state) significantly increased between E14.5 and E16.5 (5.9% ± 1.8% vs. 22.5% ± 3.3%, mean ± SD) (Figure S2D). A BrdU incorporation assay with mouse embryos also showed a substantial decrease in the ratio of proliferative cells from E14.5 to E16.5 (54.0% ± 2.1% vs. 14.3% ± 3.6%, mean ± SD) (Figures 3C and 3D). Additionally, p63^+^ cells showed a faster decrease in BrdU incorporation than p63^-^ cells (Figure S2E). These results suggest that cell cycle slowdown preferentially occurs in p63^+^ progenitors before the complete commitment into basal cell lineage and subsequently induces Krt17 expression. Based on these results, we hypothesize that epithelial quiescence occurs before basal cell specification.

**Figure 3.**
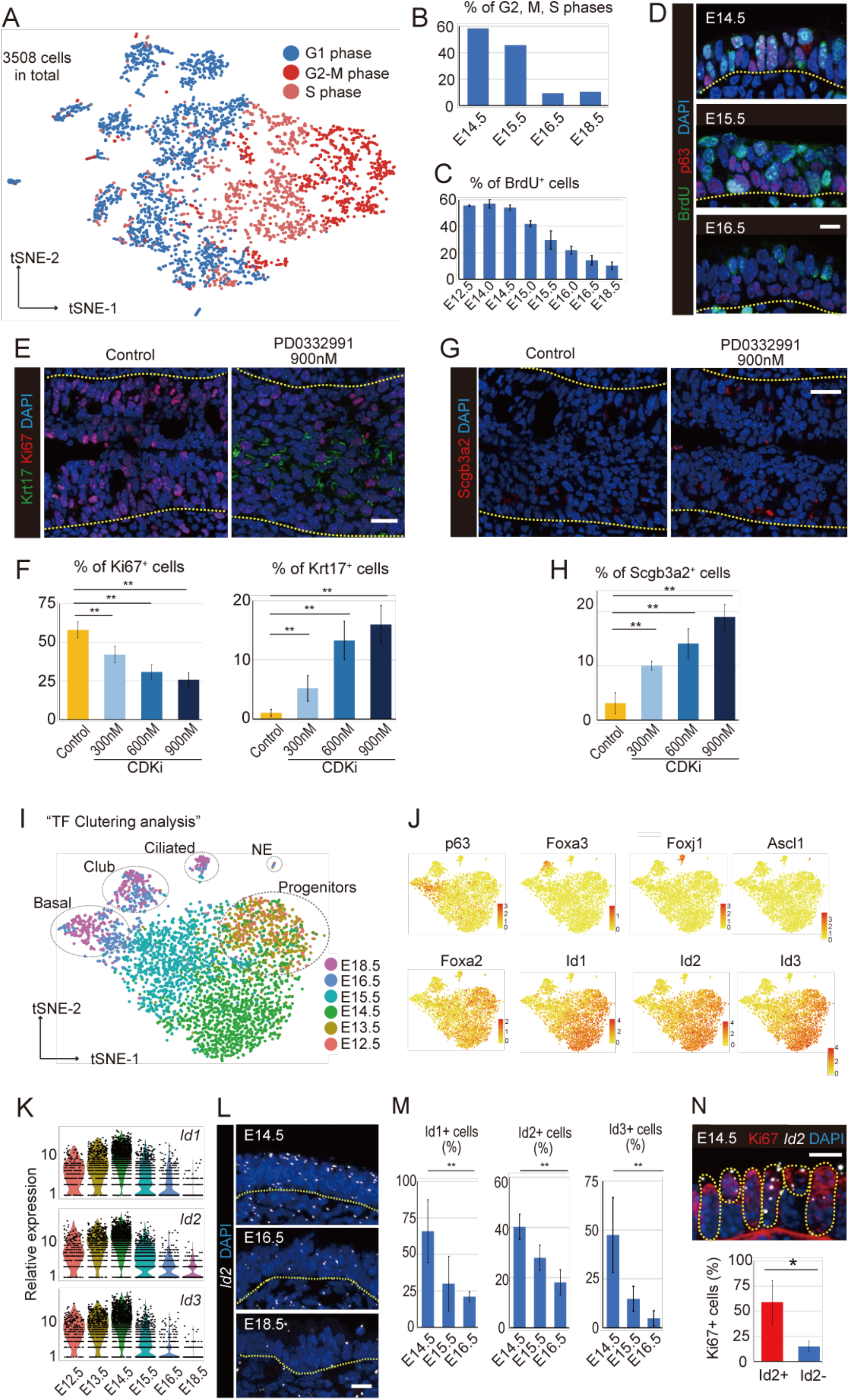
Cell cycle slowdown induces basal cell specification concomitant with downregulation of *Id* gene expression. (A) Cell cycle analysis with the scRNA-seq data using the Seurat package. (B) The ratio of cells expressing markers of S, G2, and M phases revealed a substantial decrease in proliferating cells from E14.5. (C-D) BrdU incorporation assay with immunostaining confirmed the cell cycle deceleration from E14.5. (E-G) Immunostaining for Krt17, Scgb3a2, and Ki67 in *ex vivo* cultured fetal tracheal epithelium. (F-H) Quantification of the marker-positive cells determined that PD0332991 (Cdk4/6 inhibitor) treatment induces differentiation while inhibiting proliferation in a dose-dependent manner. Reclustering analysis with TFs only (I) and expression patterns of the marker TFs (J). *Id* gene expression in the scRNA-seq data (K), PLISH for *Id2* (L), and quantification of *Id1*-, *2*-, and *3*-positive cells in PLISH (M) suggested that *Id* gene expression peaked at E14.5 and gradually decreased after that. (N) Double staining for Ki67 protein and *Id2* mRNA with immunostaining and PLISH suggested the dominant expression of Id2 in proliferating cells. * p< 0.05; Student’s t test. ** p< 0.05; Tukey’s test. Mean±SD, n=4–6 (C, M, N), 3 independent experiments (F, H). Scale bars: 5 μm (D, L, N), 10 μm (E, G).

We tested this hypothesis with *ex vivo* culture of E12.5 developing trachea, where epithelial cells are highly proliferative and do not show any lineage commitment, with and without PD0332991 (Cdk4/6 inhibitor, hereafter Cdk4/6i). Cdk4/6i inhibitor treatment induces cell cycle arrest at the G1 phase and the differentiation into *Krt17^+^* cells in a dose-dependent manner (Figures 3E and 3F). Furthermore, Scgb3a2 cells were significantly increased following Cdk4/6i treatment (Figures 3G and 3H). Hence, cell cycle arrest in epithelial progenitors is sufficient to induce basal cell specification and promote differentiation of Krt17^+^ basal progenitors.

To identify transcription factors (TFs) related to the epithelial cell cycle slowdown, we repeated cluster analysis using all 1385 genes, which are classified as TFs in Fantom5 SSTAR database (Figures 3I and 3J) (Abugessaisa, et al., 2016). The cells showed similar TF profiles at E12.5 and E13.5 but dynamically changed their TF profiles between E14.5 and E16.5, establishing distinct lineage-specific profiles. We focused on genes with expression levels that substantially changed at E14.5 and noted inhibitors of DNA-binding/differentiation genes (*Id1, Id2*, and *Id3*) (Figure S2F). Generally, Id proteins promote proliferation by antagonizing negative cell cycle regulators, such as Rb, and simultaneously inhibit differentiation by binding with various bHLH-type TFs(Roschger and Cabrele, 2017; Lasorella, et al., 2014). *Id4* was not detected in our dataset, yet the expression of *Id1, Id2*, and *Id3* was abundant in progenitors until E14.5 and then decreased along with the slowing of the cell cycle (Figure 3K). This temporal reduction in expression of *Id* genes was confirmed with proximity ligation *in situ* hybridization (PLISH)(Nagendran, et al., 2018) and dot quantification (Figures 3L and 3M). Ki67^+^ proliferative cells preferentially express *Id2* gene (Figure 3N) and *Id* genes expression was lower in Krt17^+^ cells than Krt17^-^ cells at E14.5 (Figure S2G). These observations are consistent with the hypothesis that the reduction in *Id* expression is involved in cell cycle slowdown in tracheal epithelium, which triggers the basal cell specification.

### *Id2* downregulation promotes basal cell specification of epithelial progenitors through cell cycle modulation

Next, to confirm the roles of *Id* genes in modulating epithelial cell cycle and basal cell specification, we used *Id2* loss- and gain-of-function transgenic mice. *Id2^CreERT2/CreERT2^* mice were used as *Id2* knockout mice (*Id2* KO mice)(Rawlins, et al., 2009) (Figures S3C and S3D). BrdU incorporation assays with *Id2* KO mice showed a significant decrease in proliferating cells at E14.5 and E15.5 (Figures 4A and S3A). Additionally, both Krt17^+^ and Scgb3a2^+^ progenitors were detected earlier in *Id2* KO mice than in control mice (Figure 4B). Thus, the loss of *Id2* appears to accelerate airway progenitor specification by lowering the proliferation rate. *Shh^Cre^, Rosa^3xHA-Id2-IRES-H2B-EGFP^* mice (*Id2* OE) that overexpress *Id2* in endodermal epithelial cells (Figures S3E and S3F) significantly increased *Ki67^+^* proliferative cells at E14.5 and E16.5 (Figures 4C and S3B) and decreased both *Krt17^+^* and *Scgb3a2^+^* progenitors at E14.5 (Figures 4C and 4D). Based on these observations, we conclude that the temporal regulation of Id2 dosage determines the timing of the epithelial cell cycle and basal cell specification.

**Figure 4.**
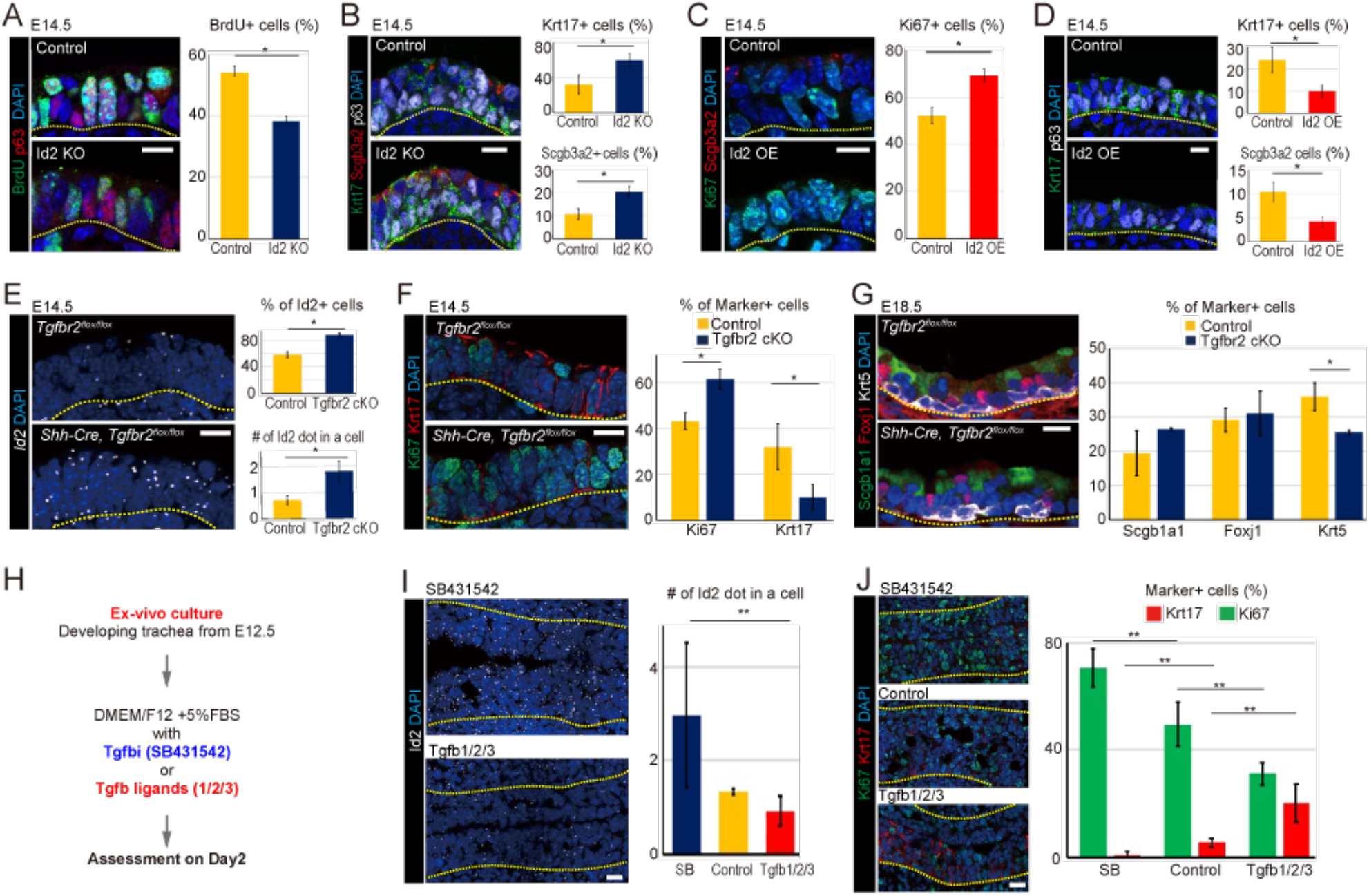
Tgfß signaling initiates cell cycle slowdown in epithelial cells by suppressing *Id2* gene expression. The phenotypic analyses of tracheas from the *Id2* KO and OE groups assessed at E14.5 by immunostaining for BrdU and p63 (A), Scgb3a2 and Krt17 (B), Ki67 and Scgb3a2 (C), and p63 and Krt17(D) revealed that *Id2* promotes proliferation while inhibiting differentiation. (E) Differences in *Id2* expression between tracheas from the control and *Tgfbr2* KO groups assessed at E14.5 with PLISH confirmed that Tgfß signaling inhibits *Id2* expression. (F) Immunostaining for Ki67 and Krt17 in *Tgfbr2* KO epithelium at E14.5 showed that loss of Tgfß signaling suppresses Krt17 expression, enhancing proliferation. (G) At E18.5, epithelial *Tgfbr2* KO resulted in a significant decrease in mature basal cells but not in other cell types. (H) Schematic representation of *ex vivo* fetal trachea culture. Expression of *Id2* mRNA (PLISH) (I) and Ki67/Krt17 protein (immunostaining) (J) in *ex vivo* cultured trachea after 2 days of treatment with Tgfß-1/2/3 ligands or SB431542 confirmed that Tgfß signaling suppresses *Id2* and Ki67 expression, increasing Krt17 expression. * p< 0.05; Student’s t test. ** p< 0.05; Tukey’s test. Mean±SD, n=4-6 (A-G), 3 independent experiments (I-J). Scale bars: 5 μm.

### Mesenchymal-to-epithelial Tgfß signaling initiates cell cycle slowdown in epithelial cells by suppressing the *Id2* gene

Since *Id* genes are known to be downstream factors of Tgfβ signaling in hematopoietic cells (Roschger and Cabrele, 2017; Lasorella, et al., 2014), we evaluated the effects of Tgfβ signaling on *Id* genes using *Shh^Cre^, Tgfbr2^flox//flox^* mice in which the *Tgfβ receptor 2* gene is ablated in the endodermal epithelium (*Tgfbr2* cKO). The *Id2* gene expression was significantly upregulated in the *Tgfbr2* cKO mice at E14.5 (Figure 4E), suggesting that Tgfβ inhibits *Id2* expression. Consistent with the enhanced *Id2* phenotype, epithelial cells in *Tgfbr2* cKO mice showed more *Ki67^+^* proliferative cells and a more delayed appearance of *Krt17^+^* progenitors than those in the control mice (Figure 4F). At E18.5, *Tgfbr2* cKO mice showed a significant decrease in mature *Krt5*^+^ basal cells (Figure 4G), indicating that *Tgfbr2* activation is involved in basal cell specification by inhibiting *Id2* expression. These findings are consistent with the previous report that Smad activation through Tgfβ signaling is necessary for the specification of human *p63^+^* basal cell population from *Id2^+^* primordial progenitors (Miller, et al., 2020). *Ex vivo* trachea culture with Tgfβ ligands (Tgfβ-1/2/3) confirms that Tgfβ ligand inhibits *Id2* expression, forces epithelial cells into slow cycling, and induces more *Krt17*^+^ cells than Tgfβ inhibitor (SB431542) (Figures 4H-J). Hence, we propose that Tgfβ signaling acts as the initial cue for the epithelial cell cycle slowdown and the specification of the *Krt17^+^* basal progenitors by decreasing *Id2* gene expression.

Next, we investigated Tgfβ ligand-secreting cells with PLISH and found that mesenchymal cells express *Tgfβ-3* throughout development (Figure S4A). The number of Tgfβ-3-secreting cells decrease over time, but subepithelial mesenchymal cells are positive for *Tgfβ-3* even at E18.5. Thus, it is highly likely that airway mesenchyme is the major source of the Tgfβ ligand.

### *Id2* expression in mature basal stem cells ensures their proliferative potential at perinatal and adult stages

While we showed that *Id2* attenuation is the key step for the specification of basal cell progenitors in development, the predominant expression of *Id* genes in mature basal cells has been reported in adulthood (Montoro et al., 2019). We compared *Id2* expression between basal and luminal cells isolated from Krt17-EGFP transgenic mice (Bianchi, et al., 2005) at E18.5 using qRT-PCR and confirmed twofold higher expression of *Id2* in basal cells in compare to luminal cells (Figure 5A). Therefore, next, we asked whether the *Id2* gene’s function in regulating proliferation during development was conserved at perinatal and adult stages. We employed a tracheosphere culture assay and estimated the colony-forming efficiency (CFE) in *Id2* KO, control, and *Id2* OE mice at E18.5 (Figure 5B). The epithelial cells from the *Id2* OE mice showed a significantly higher CFE than those in other groups, suggesting that the Id2 dosage was related to the proliferative capacity of epithelial cells at the perinatal stage. The predominant expression of *Id2* in basal cells was also detected in 2-month-old adult mice (Figure 5A). Similar to E18.5, basal cells from *Id2* OE mice showed a significantly higher CFE than those in the control group (Figure 5C). These results confirm that the Id2 dosage ensures the self-renewal capacity of basal cells at perinatal and adult stages.

**Figure 5.**
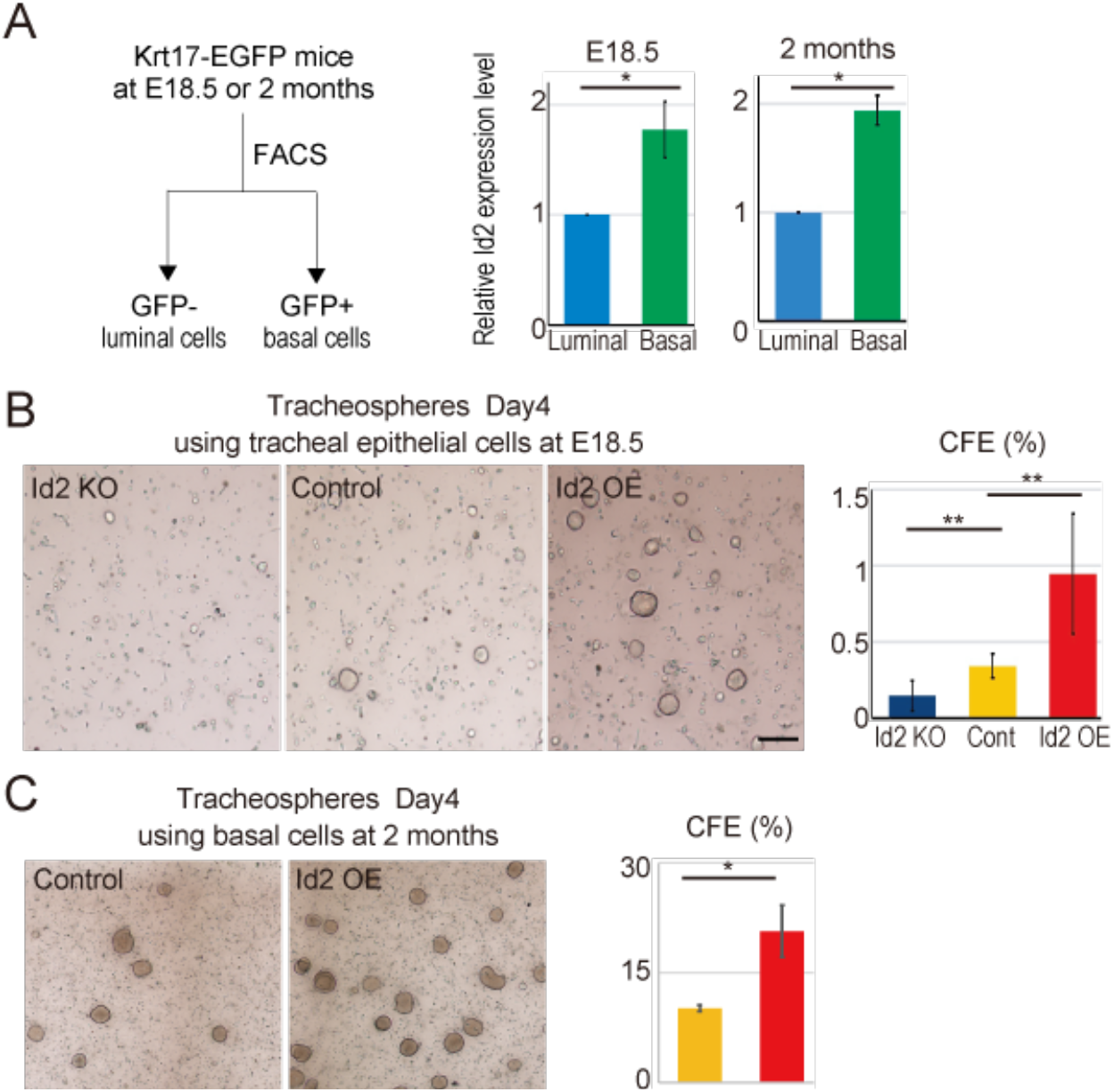
Predominant *Id2* expression in mature basal cells ensures their proliferative potential at perinatal and adult stages. (A) Krt17^+^ and Krt17^-^ cells were sorted based on GFP intensity from the tracheas of Krt17-EGFP mice at E18.5 or 2 months, and *Id2* expression was assessed with qRT-PCR. *Id2* expression was significantly higher in the Krt17^+^ cells than in the Krt17^-^ cells at both timepoints. Tracheosphere culture at day 4 using tracheal epithelial cells derived from the *Id2* KO, control, and *Id2* OE mice at E18.5 (B) or 2 months (C) confirmed that the CFE reflects the dosage of *Id2* at both timepoints. * p< 0.05; Student’s t test. ** p< 0.05; Tukey’s test. Mean±SD, 4 independent experiments (A-C). Scale bars; 300 μm.

### The slow-cycling basal cells re-enter the cell cycle with recurrent *Id2* activation following injury

Next, we asked whether *Id2* was involved in airway tissue regeneration at the adult stage. SO_2_-inhalation mediated airway injury can be used as a model to study basal cell-driven airway epithelial regeneration (Pardo-Saganta, et al., 2015; Rawlins, et al., 2007; Borthwick, et al., 2001). Therefore, we used this model to assess the airway epithelial regeneration process. The number of Ki67-expressing cells started increasing at 18 h post-injury (hpi), and peaked at approximately 24 hpi (Figure 6A), suggesting that the transition of basal cells from slow-cycling to active proliferation occurs around 18 hpi. An increase in *Id2* expression was detected at 12 hpi before the increase in Ki67-expressing cells (Figure 6B). These observations prompted us to hypothesize that recurrent Id2 activation stimulated basal cells to re-enter the active cell cycle. We exposed wild-type and *Id2* OE adult mice to SO_2_ gas and examined their respective number of Ki67-expressing cells to test this hypothesis. Airways from the *Id2* OE mice showed a more rapid response to injury than airways from wild-type mice at 12 hpi (Figure 6C), indicating the artificially increased *Id2* expression enhances basal cell proliferation. Additionally, *Id2* OE prolonged the proliferation of Krt5^+^ basal cells more than 72 hpi and prohibited basal cell differentiation into Krt5^-^ luminal cells, showing the phenotype similar to basal cell hyperplasia, a precancer-like state (Figure 6C). These results imply that high *Id2* expression in airway epithelial cells promoted the transition of basal cells from the slow-cycling to the active proliferation state; in contrast, sustained *Id2* expression inhibits a return from the active proliferation state to the slow-cycling state. Thus, *Id2* is a key factor in regulating basal cells’ transition from slow cycling to proliferation in response to epithelial damage and inhibiting their differentiation into luminal cells.

**Figure 6.**
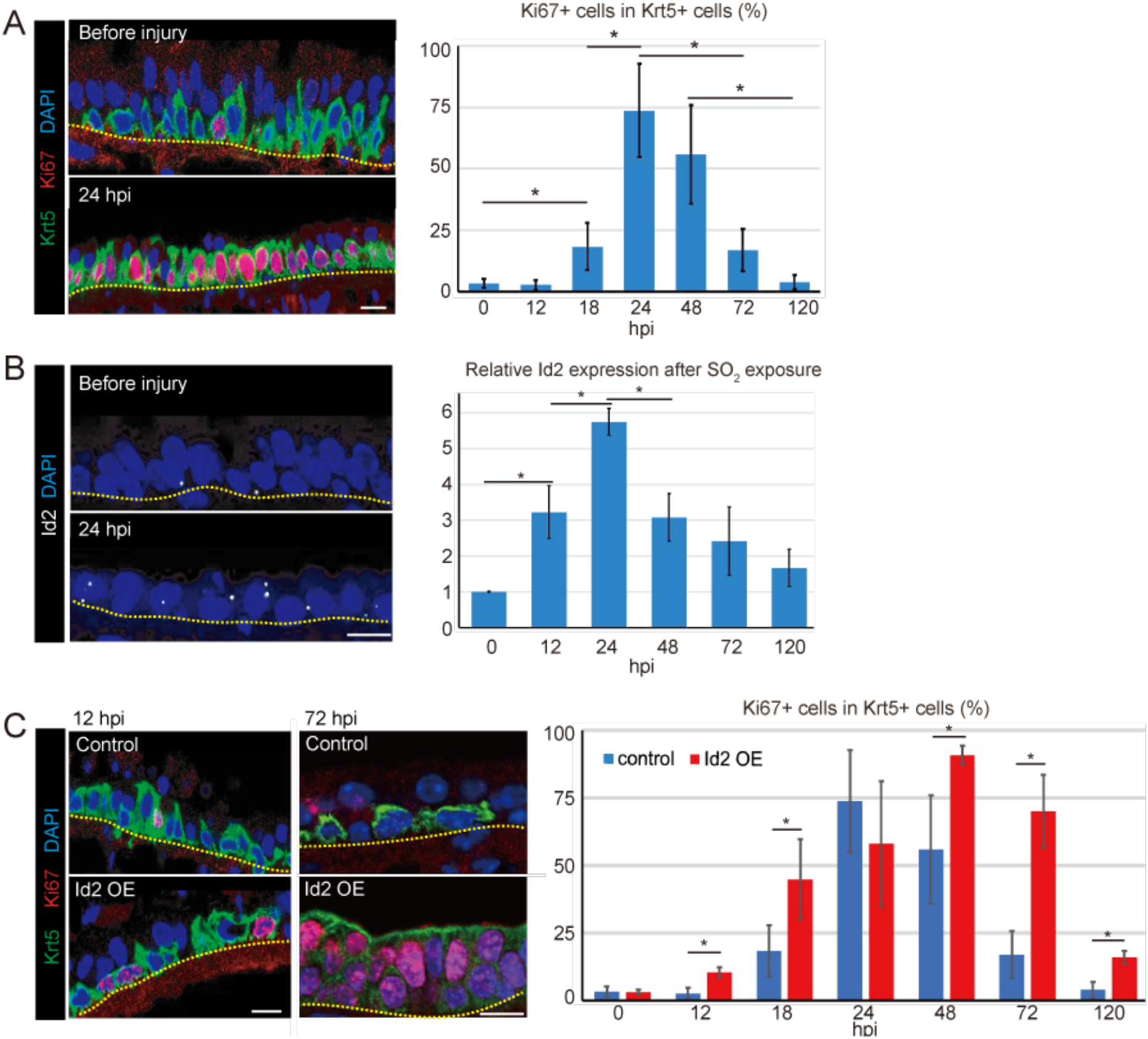
The slow-cycling basal cells re-enter the cell cycle with recurrent *Id2* activation following SO_2_ injury. (A) Assessment of proliferating basal cells in injured airway epithelium by immunostaining for Ki67 and Krt5. Ki67^+^ proliferative cells in the Krt5^+^ basal cell population started increasing at 18 hpi and peaked at 24 hpi in adult mice. (B) *Id2* expression during regeneration was assessed by PLISH. *Id2* expression started increasing at 12 hpi prior to the increase in Ki67^+^ cells and peaked at 24 hpi. (C) Ki67 expression pattern assessed by immunostaining in the control and Id2 OE mice before or after SO_2_ exposure confirmed that *Id2* dosage promotes basal cell self-renewal during the repair process. hpi, hours post injury. * p< 0.05; Tukey’s test. Mean±SD, n=4-6 (A-C). Scale bars: 5 μm.

### The inhibitory effect of Tgfβ signaling on Id2 expression is conserved until adult stages

Lastly, we attempted to determine that the inhibitory effect of Tgfβ signaling on Id2 expression in lung development is conserved at perinatal and adult stages. First, we checked the expression pattern of pSmad2/3 at E18.5, which is downstream of Tgfβ signaling. Consistent with predominant *Id2* expression in mature basal cell population, pSmad2/3 expression is significantly lower in mature basal cells than luminal cells (Figure 7A). In addition, Tgfβ superfamily signaling inhibitors, such as *Tgif1, Nbl1, Sostdc1*, and *Fst* are highly and exclusively expressed in basal cell population (Figures S5A-F). These observations suggest that basal cells may express Tgfβ inhibitors to inhibit Tgfβ signaling and maintain *Id2* expression. Supporting this idea, dual Smad inhibition via Tgfβ/Bmp inhibitors is beneficial for the maintenance of mature basal cells (Mou, et al., 2016; Tadokoro, et al., 2016). The inhibitory effect of Tgfβ signaling on *Id2* expression was directly confirmed by the basal cell culture with Tgfβ ligands (Tgfβ-1/2/3) or inhibitor (SB431542) (Figure 7B). Tgfβ ligands treatment significantly decreased *Id2* expression in basal cell culture.

**Figure 7.**
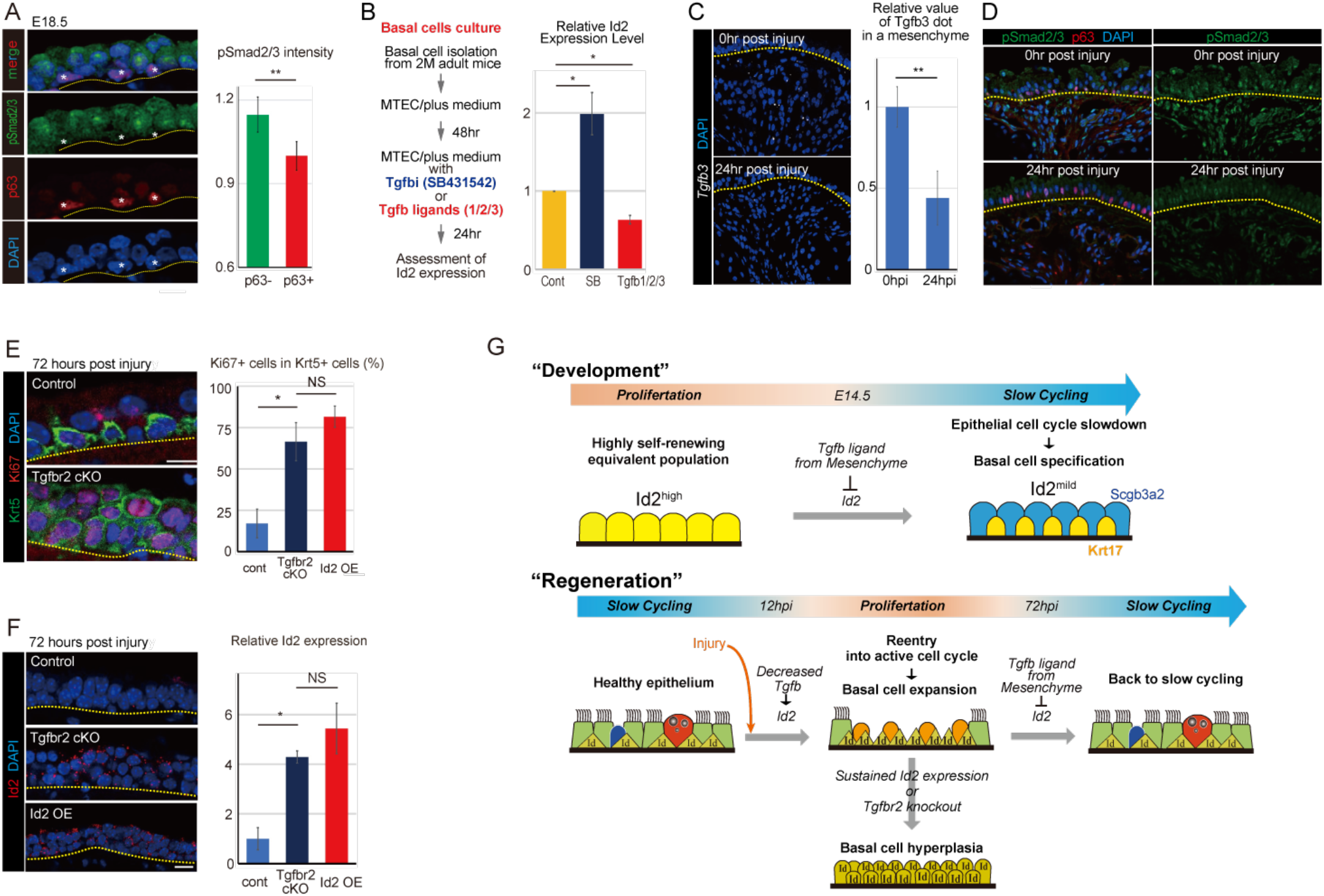
The inhibitory effect of Tgfβ signaling on *Id2* expression is conserved until adult stage. (A) Quantitative assessment of pSmad2/3 staining intensity showed higher expression in the p63^-^ luminal cells than in the p63^+^ basal cells. Asterisks indicate the p63^+^ basal cells. (B) Basal cells were sorted based on EpCAM/NGFR from the tracheas of 2-month-old wild-type mice, and *Id2* expression was assessed with qRT-PCR after 1 day culture with Tgfb ligands(1/2/3 or Tgfb inhibitor(SB431542). Tgfb inhibitor treatment significantly increased *Id2* expression, while Tgfb ligands treatment significantly decreased *Id2* expression. (C) *Tgfb3* detected by PLISH at 0 and 24 hpi confirmed the significant decrease of Tgfb3 expression in mesenchymal cells after SO_2_ injury. (D) Immunostaining with anti-pSmad2/3 antibody demonstrated the decreased expression of pSmad2/3 in both epithelial and mesenchymal cells after SO_2_ injury. (E) Assessment of proliferating basal cells in injured airway epithelium by immunostaining for Ki67 and Krt5. Tgfbr2 cKO mice phenocopied the basal cell hyperplasia phenotype seen in *Id2* OE mice. (F) *Id2* expression at 72hpi was assessed with RNAscope. *Id2* expression was significantly increased in *Tgfbr2* cKO and *Id2* OE mice compared to control. (G) Schematic summary of slow-cycling basal cell specification during development. *Id2* attenuation triggered by Mesenchymal-to-epithelial Tgfβ signaling slows down the cell cycle and contributes to the specification of Krt17^+^ basal cell progenitors. See discussion for the details. Schematic summary of the function of *Id2* gene during tissue regeneration following SO_2_ injury. SO_2_ injury decreases Tgfb3 secretion from mesenchyme, which reactivates *Id2* expression in basal cells. Recurrent *Id2* activation initiates basal cell expansion, but its sustained expression results in basal cell hyperplasia. See discussion for the details. hpi, hours post injury. * p< 0.05; Tukey’s test. ** p< 0.05; Student’s t test. NS; Not significant. Mean±SD, n=4-6 (A-C, E, F). Scale bars: 5 μm.

We further assess the function of Tgfβ signaling in *in vivo* tissue regeneration process. *Tgfβ–3* and pSmad2/3 expression after SO_2_ injury was monitored with PLISH and immunostaining, respectively. *Tgfβ–3* expression was significantly decreased after SO_2_ injury (Figures 7C and 7D), consistent with the decreased pSmad2/3 intensity in both epithelial and mesenchymal cells. Furthermore, *Tgfbr2* cKO mice at 72 hpi also showed increased *Id2* expression (Figure 7F) as well as basal cell hyperplasia, the phenocopy of *Id2* OE mice (Figure 7E). Thus, the Tgfβ-Id2 axis is likely a critical regulator of the transition between the active proliferation and the slow-cycling state, which is conserved during development and adult tissue regeneration, in airway stem cells.

## Discussion

In the present study, we investigated the conserved mechanism regulating the proliferation mode transitions of airway basal stem cells in development and regeneration; we also found that Tgfβ-Id2 axis is a commonly shared regulator in both these processes. During basal cell specification in airway development, *Id2* attenuation triggered by mesenchymal-to-epithelial Tgfβ signaling slows down epithelial progenitors’ cell cycle and induces the *Krt17^+^* basal progenitors. *Id2* expression ensures mature basal cells’ capacity for self-renewal in a dose-dependent manner. In the adult tissue regeneration model, the recurrent activation of *Id2* via Tgfβ reduction initiates tissue regeneration by forcing the slow-cycling basal cell to re-enter the active cell cycle. While proliferating basal cells get back to the slow-cycling state by 120 hpi in normal tissue regeneration, enhanced *Id2* expression or impaired Tgfβ receptor results in basal cell hyperplasia that resembles a precancerous condition (Figure 7G).

Basal cells’ tightly regulated proliferative potential is critical for tissue regeneration, homeostasis, and the avoidance of pathological conditions. Airway basal cells remain quiescent under homeostasis; in response to injury, they re-enter the cell cycle to replenish the lost cells by producing transit-amplifying cells. However, excessive proliferation is related to squamous cell carcinoma (Lapouge, et al., 2011). Recently, there has been increasing evidence of tumor cells, including lung cancer cells, hijacking the embryonic pathways, which control stem and progenitor cell behavior during development (Laughney, et al., 2020; Tata, et al., 2018; Murry and Keller, 2008). Following this concept, we demonstrated that mature basal stem cells reutilized the Tgfβ-Id2 axis in tissue regeneration for the tightly regulated transitions between slow-cycling and proliferation. During development, the Tgfβ-Id2 axis also plays critical roles in regulating the cell cycle state for the specification of slow-cycling basal progenitors.

The acquisition of cellular quiescence in stem cell development has been reported in mammalian neurogenesis (Furutachi, et al., 2015). A subset of embryonic neuroepithelial cells slows cell cycling between E13.5 and E15.5, isolating a population of primordial neural stem cells. The expression of the cyclin-dependent kinase inhibitor, p57, demarcates primordial neural stem cells from non-stem cells and defines reversible quiescence status and specification numbers of adult neural stem cells. The embryonic origin of adult stem cells could acquire quiescence by reducing the number of divisions at an early stage of development to prevent the exhaustion of the stem cell pool. (Cheung and Rando, 2013).

Id2 was first reported as a marker of multipotent cells at the distal tip region of developing lungs(Rawlins, et al., 2009). This region includes highly self-renewing progenitors. However, Id2 function in airway stem cell is still to be determined. We show that *Id2* is highly expressed in primordial progenitors in developing airways until E14.5 (Figures 3J-M). Generally, *Id* genes display two direct effects on proliferation and differentiation via independently interacting with negative cell cycle regulators, such as Rb, and differentiation-related TFs, such as bHLH-type TFs(Roschger and Cabrele, 2017; Lasorella, et al., 2014). *Id* gene dosage during airway development could determine the proliferation state because *Id2* loss-of-function transgenic mice show advanced cell cycle attenuation and basal cell specification. In contrast, these processes were delayed in *Id2* gain-of-function transgenic mice (Figures 4A and 4C). We cannot exclude the possibility that Id2 directly inhibits key TFs for epithelial differentiation, although we did not detect any bHLH-type TFs as promising candidates in our scRNA-seq data. In future studies, a comprehensive screening assay is required to confirm the existence of specific TFs responsible for epithelial differentiation.

TGF superfamily signaling is reported to induce stem cell quiescence in various organs(Genander, et al., 2014; Kandasamy, et al., 2014; Nishimura, et al., 2010; Yamazaki, et al., 2009). In the present study, we showed that Tgfβ signaling initiates the transition in airway epithelial cells from the active proliferation to the slow-cycling state by suppressing *Id2* expression; this transition is most likely conserved during both development and regeneration. Generally, Tgfβ activates or suppresses *Id* genes context-dependently(Roschger and Cabrele, 2017; Lasorella, et al., 2014). Tgfβ inhibits *Id* genes in basal keratinocytes that are similar to airway basal cells(Rotzer, et al., 2006). This inhibitory effect of Tgfβ is also consistent with dual inhibition of Tgfβ/BMP signaling that promotes self-renewal of the basal cell population(Mou, et al., 2016; Tadokoro, et al., 2016). However, the precise molecular mechanism how Tgfβ signaling inhibits *Id2* expression is still to be elucidated.

The present study shows that mature basal cells maintain moderate Id2 expression to ensure proliferative potential at perinatal and adult stages (Figure 5A). Our organoid culture experiments confirmed the positive function of *Id2* in epithelial proliferation (Figures 5B and 5C), indicating that *Id2* has a consistent function in basal cell lineage to promote proliferation in a dose-dependent manner (Figure 7G). This critical role of Id genes is conserved in neural and hematopoietic stem cells(Jung, et al., 2010; Jankovic, et al., 2007). Additionally, Id2 dosage directly affects the normal repair processes in response to SO_2_ injury (Figure 6C). After SO_2_ injury, the artificial enhancement of the *Id2* gene accelerates tissue regeneration, but aberrant *Id2* gene expression results in premalignant basal cell hyperplasia seen in *Id2* OE mice. Thus, the tight regulation of Id2 is needed for normal tissue regeneration prohibiting pathological conditions due to sustained proliferation. Consistent with this conclusion, aberrant Id genes expression is observed in cancer cells in various organs, especially in cancer stem cells including the lungs, contributing to the tumorigenesis and metastasis (Roschger and Cabrele, 2017; Lasorella, et al., 2014; Pillai, et al., 2011).

In addition, the fact that Tgfbr2 cKO mice phenocopied Id2 OE mice after SO_2_ injury demonstrated that Tgfβ signaling governs tissue regeneration via controlling proliferative states of basal stem cells through tight regulation of Id2 expression (Figures 7E and 7F). Thus, fine tuning of Tgfβ-Id2 axis is a key for proper recovery from severe tissue damage caused by influenza virus or SARS-CoV-2 and represents a possible therapeutic target in squamous lung carcinoma.

## Acknowledgments

We thank David M. Owens for the Krt17-CreER mice, Tasuku Honjo for the Rbpj flox mice, Raphael Kopan for the N2IP::Cre mice and the Animal Resource Development Unit. We also thank Kuraku Shigehiro, Kadota Mitsuru, Nishimura Osamu, Quan Wu, and Miura Hisashi for assistance with the scRNA-seq data analysis. We thank Raphael Kopan and Hiroshi Hamada for reviewing the manuscript.

These studies are supported by funding from Grants-in-Aid for Scientific Research (B) (20H03693) (M.M.), Young Scientists (19K17691) (K.H.) of the Ministry of Education, and Culture, Sports, Science and Technology, Japan; from RIKEN BDR-Otsuka Pharmaceutical Collaboration Center (RBOC) and from the Special Postdoctoral Researcher (SPDR) Program of RIKEN (H.K.).

## Author contributions

K.H. and M.M. designed the project and performed experiments with the aid of Y.A. and M.C. K.H. analyzed the single-cell transcriptomics data from embryonic and adult tracheal epithelium. Y.A. assisted with the mouse experiments. M.C. supported the generation of LSL-Id2-IRES-H2B-EGFP. A.T. and K.H. generated the Rosa26^LSL-Id2-IRES-H2B-EGFP^ animals. K.H. and M.M. wrote the manuscript with the contribution of all authors.

---

**Figure S1.**
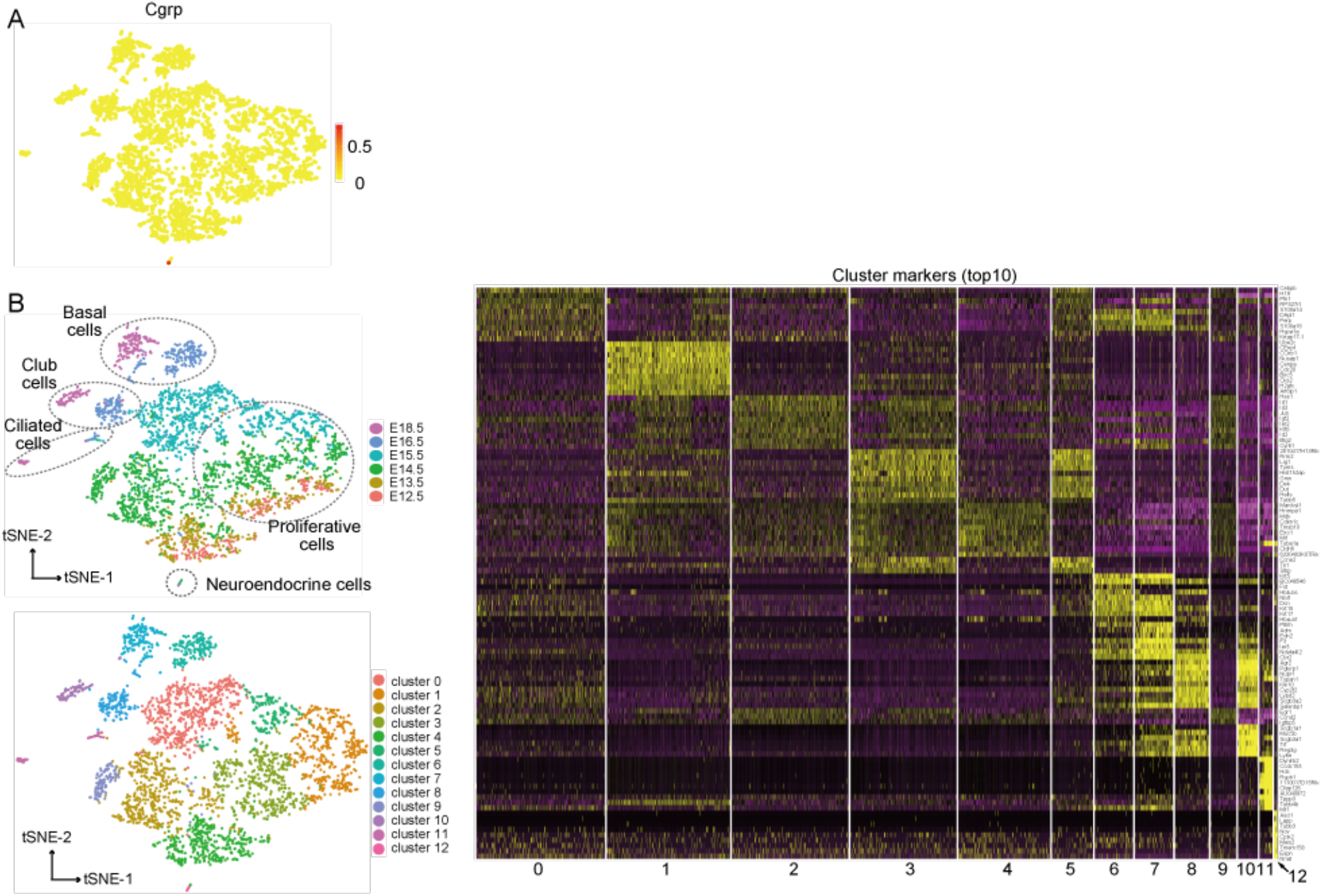
scRNA-seq analyses using embryonic progenitors. (A) Cgrp expression in the scRNA-seq dataset. (B) Clustering analysis and cluster markers (top 10) in a heatmap using embryonic progenitors from E12.5 to E18.5.

**Figure S2.**
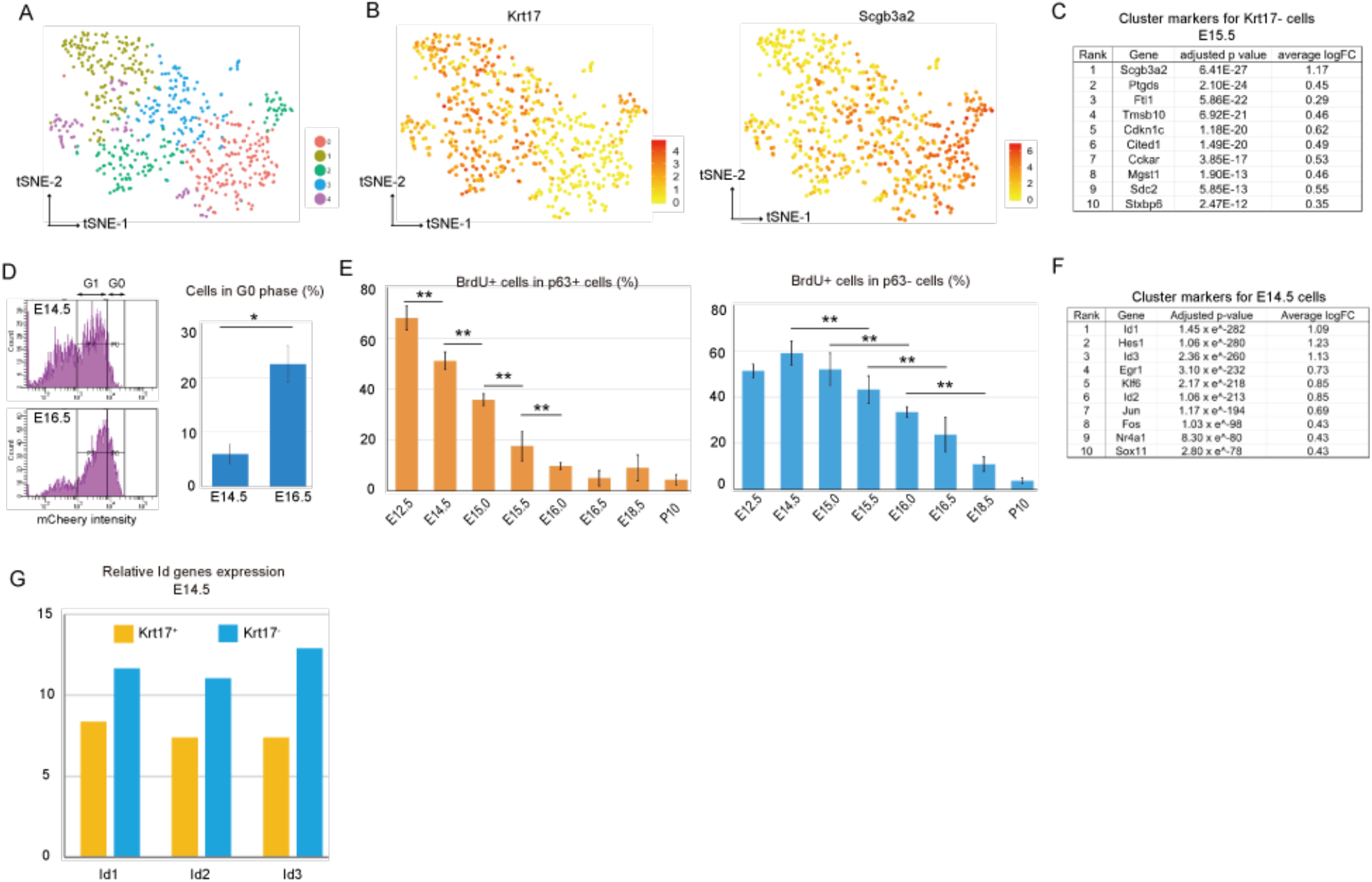
scRNA-seq analyses using E15.5 epithelial progenitors only and proliferative profile analyses. (A) Clustering analysis using epithelial cells from the E15.5 trachea only. (B) The mutually exclusive expression patterns of *Krt17* and *Scgb3a2* in E15.5 scRNA-seq data. (C) Top 10 cluster marker list for *Krt17^-^* cells at E15.5 based on the E15.5 scRNA-seq data. (D) The percentage of cells in G0 phase detected in Fucci mice (*Shh-Cre, Rosa^H2B-EGFP/FucciG1^*) significantly increased at E16.5 compared with E14.5. (E) BrdU incorporation assay with immunostaining confirmed the earlier decline of BrdU^+^ proliferative cells in p63^+^ cell population than p63^-^ cell population. (F) Top10 cluster markers for the E14.5 cluster. (G) Id genes expression, which was calculated with E14.5 scRNA-seq dataset, was decreased in Krt17^+^ cells than Krt17^-^ cells at E14.5. * p< 0.05; Student’s t test. ** p< 0.05; Tukey’s test.

**Figure S3.**
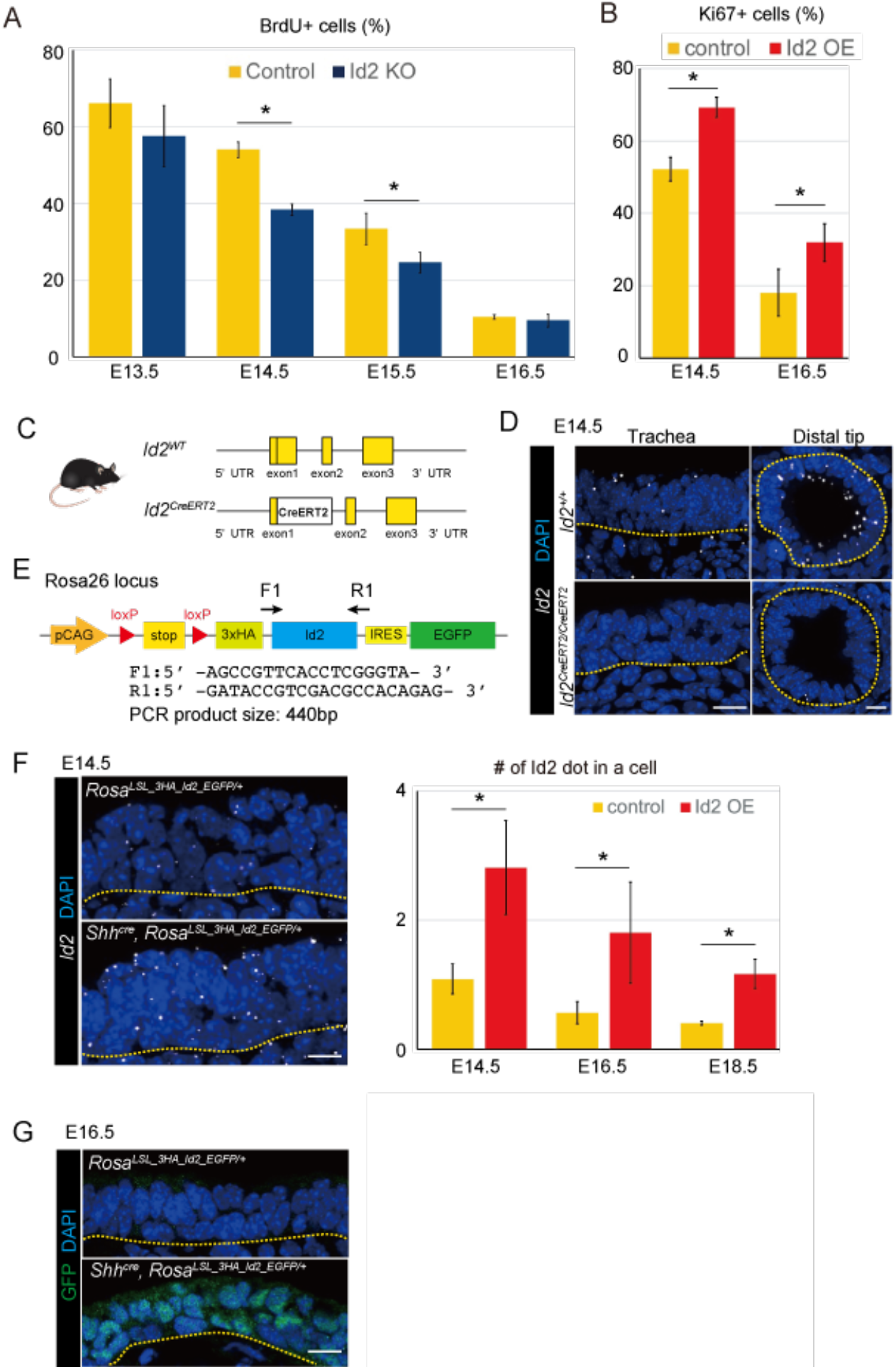
Phenotypes of *Id2* knockout and overexpression mice. (A) Time-series immunostaining analysis showed more BrdU^+^ epithelial cells in the *Id2*^+/+^ mice (control) than the *Id2^CreERT2/CreERT2^* mice (*Id2* KO mice) at E14.5 and E15.5. (B) Time-series immunostaining analysis shows fewer Ki67^+^ epithelial cells in the *Rosa^3xHA-Id2-IRES-H2B-EGFP^* mice (control) than the *Shh^Cre^, Rosa^3xHA-Id2-IRES-H2B-EGFP^* mice (*Id2* OE) at E14.5 and E16.5. (C) Schematic summary of the *Id2* locus in the wild-type and Id2^CreERT2/+^ mice. (D) *Id2* expression detected by PLISH in the trachea and distal tip epithelium at E14.5 validates the knockout of *Id2* expression in the *Id2* KO mice. (E) Inserted construct in the Rosa26 locus of the *Rosa^3xHA-Id2-IRES-H2B-EGFP^* mice. (F) PLISH image analysis of *Id2* expression confirmed the overexpression of *Id2* in the *Id2* OE mice. (G) GFP detection with immunostaining at E16.5 in the control and *Id2* OE mice. * p< 0.05; Student’s t test. Scale bars: 5 μm.

**Figure S4.**
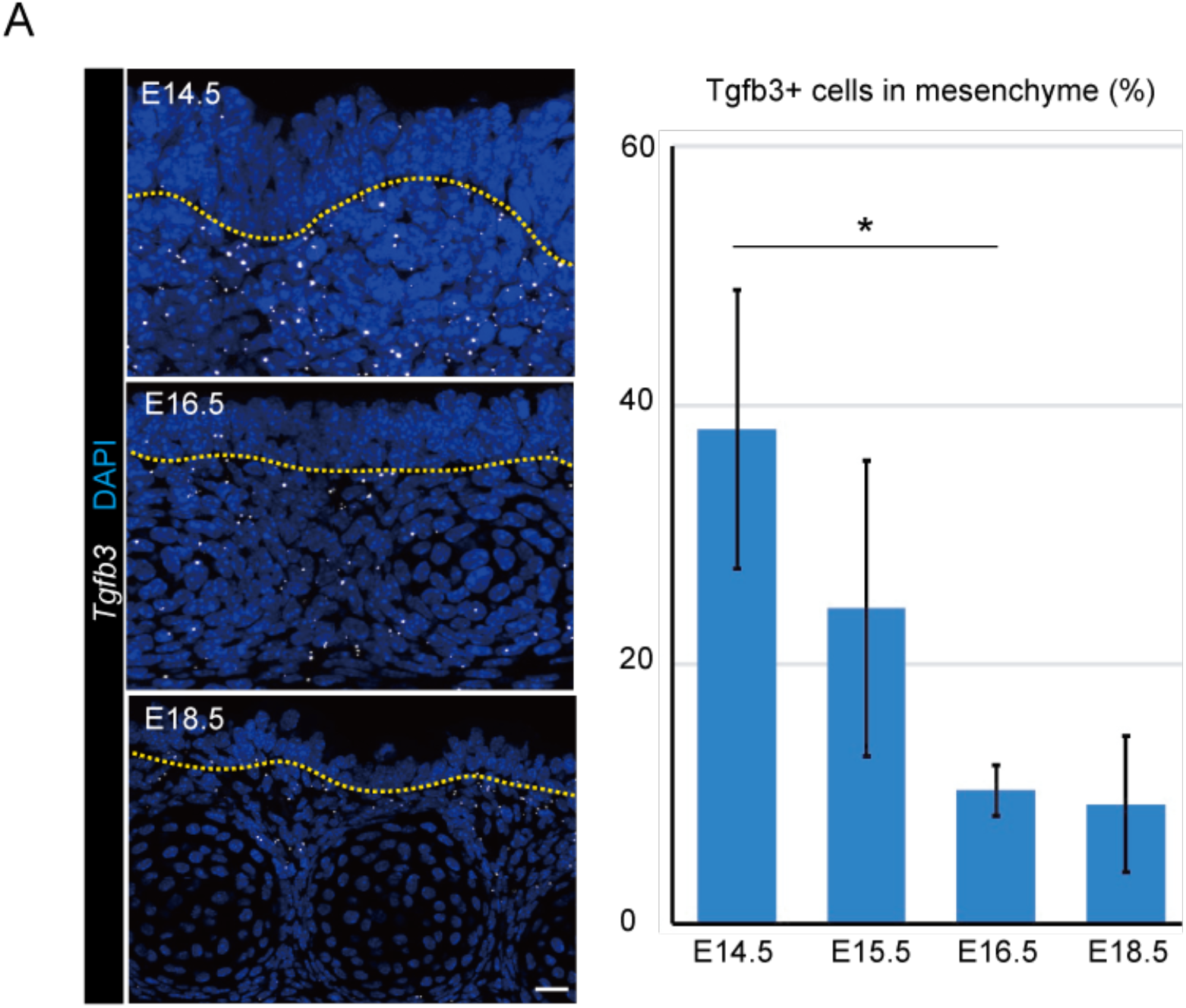
*Tgfb3* expression in the developing trachea. (A) *Tgfb3* detected by PLISH in the developing trachea from E14.5 to E18.5 confirmed that mesenchymal cells are the main source of Tgfb3 ligand during development, and their expression decreases over time. * p< 0.05; Tukey’s test. Scale bars: 10 μm.

**Figure S5.**
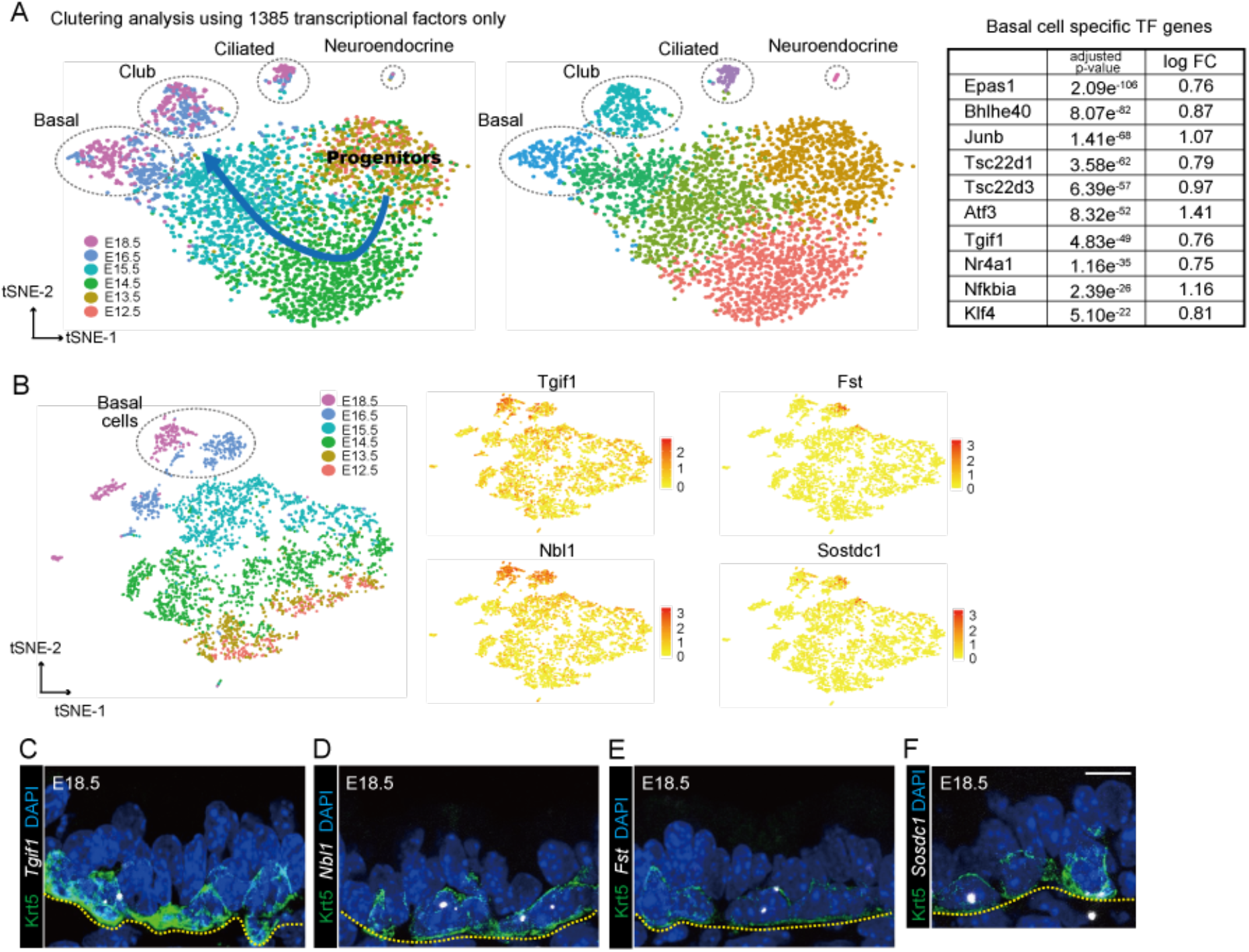
Basal cell lineage-specific genes detected in the scRNA-seq dataset. (A) The top 10 list of basal cell-specific transcription factors (TFs) detected by clustering analysis using TFs only. (B) The expression of 4 genes (*Tgif1, Nbl1, Fst*, and *Stdc1*) in the t-SNE map that are specifically expressed in basal cells in the later stage of development. The spatial expression patterns of *Tgif1* (C), *Nbl1* (D), *Fst* (E), and *Stdc1* (F) detected by PLISH. Scale bars: 5 μm.

## STAR⋆METODS

Detailed methods are provided in the online version of this paper and include the following:

● KEY RESOURCES TABLE
● LEAD CONTACT AND MATERIALS AVAILABILITY
● EXPERIMENTAL MODEL

○ MICE
● METHOD DETAILS

○ BrdU-incorporation assay
○ Cell cycle analysis
○ Cell dissociation and FACS
○ Ex-vivo trachea culture experiment
○ Microscopy and imaging
○ Immunohistochemistry
○ Quantitative RT-PCR (qPCR)
○ Single cell RNA-seq for sequencing library construction
○ Single cell RNA-seq analyses
○ Single molecule in-situ hybridization (PLISH)
○ Single molecule in-situ hybridization (RNAscope)
○ SO_2_ airway injury model
● QUANTIFICATION AND STATISTICAL ANALYSIS

○ Statistical analysis
● DATA AND CORE AVAILABILITY

○ Antibodies
○ Chemicals, Peptides, and Recombinant Proteins
○ Critical Commercial Assays
○ Deposited Data
○ Experimental Models: Organisms/Strains
○ Oligonucleotides
○ Software and Algorithms

### KEY RESOURCES TABLE

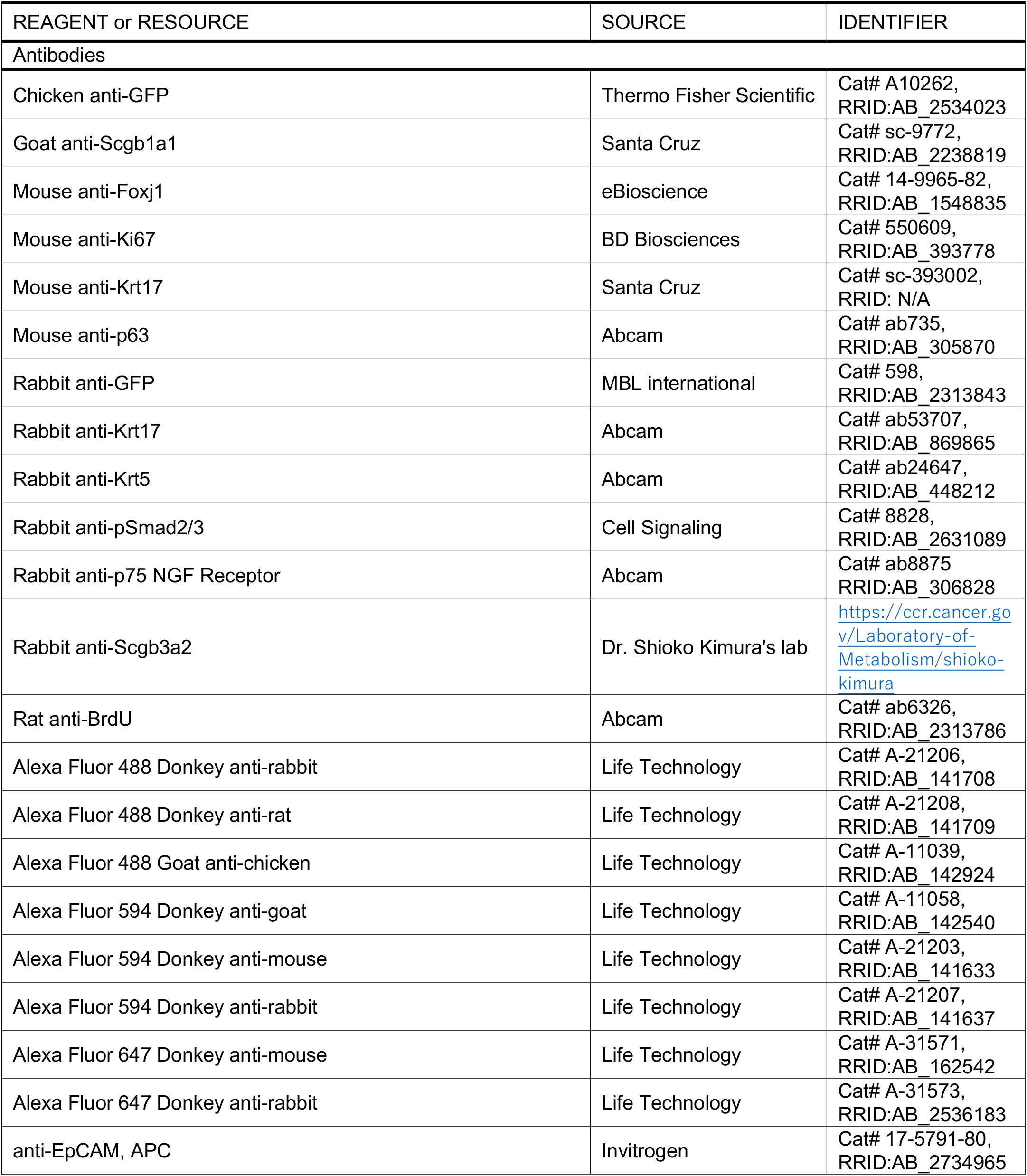
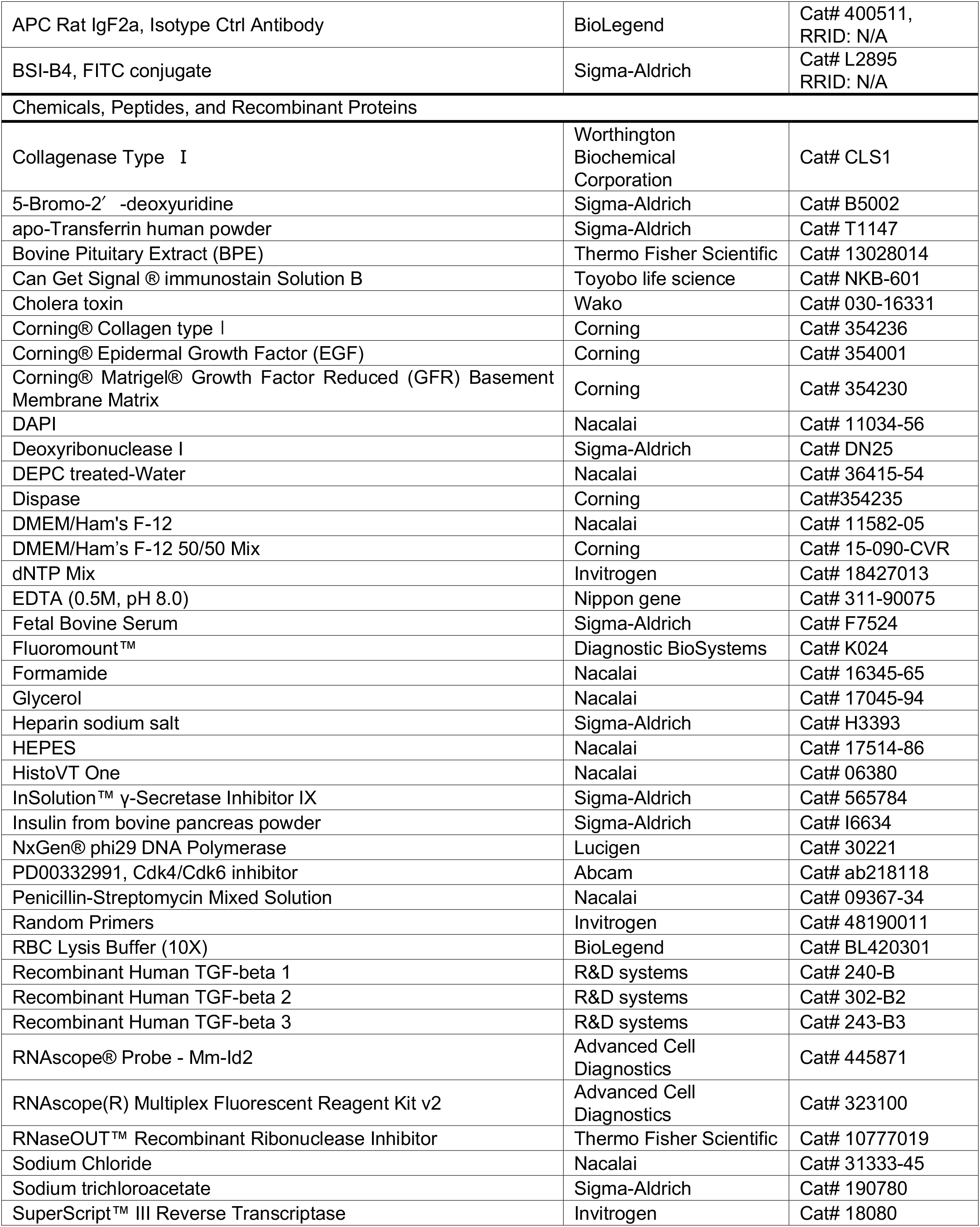
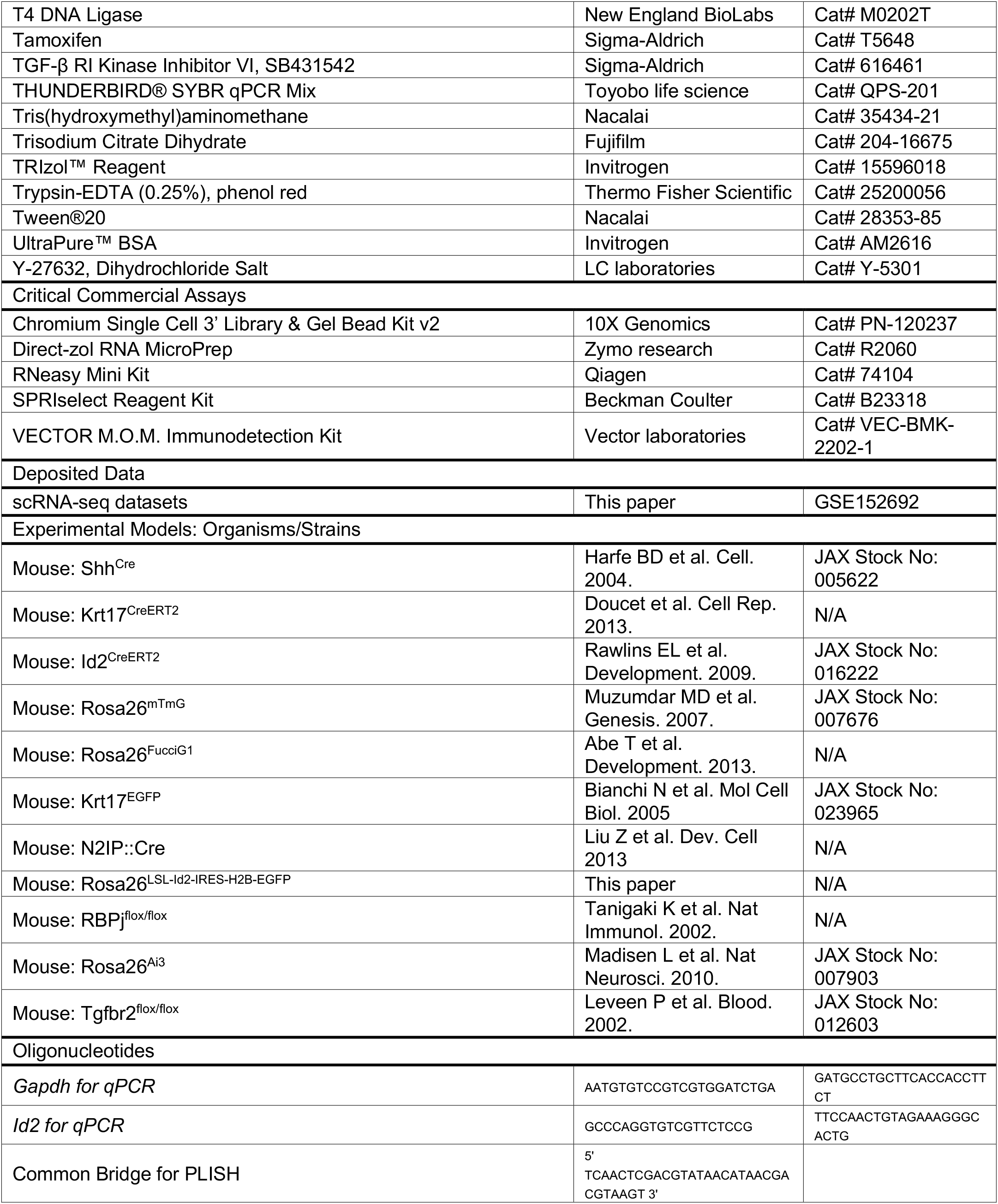
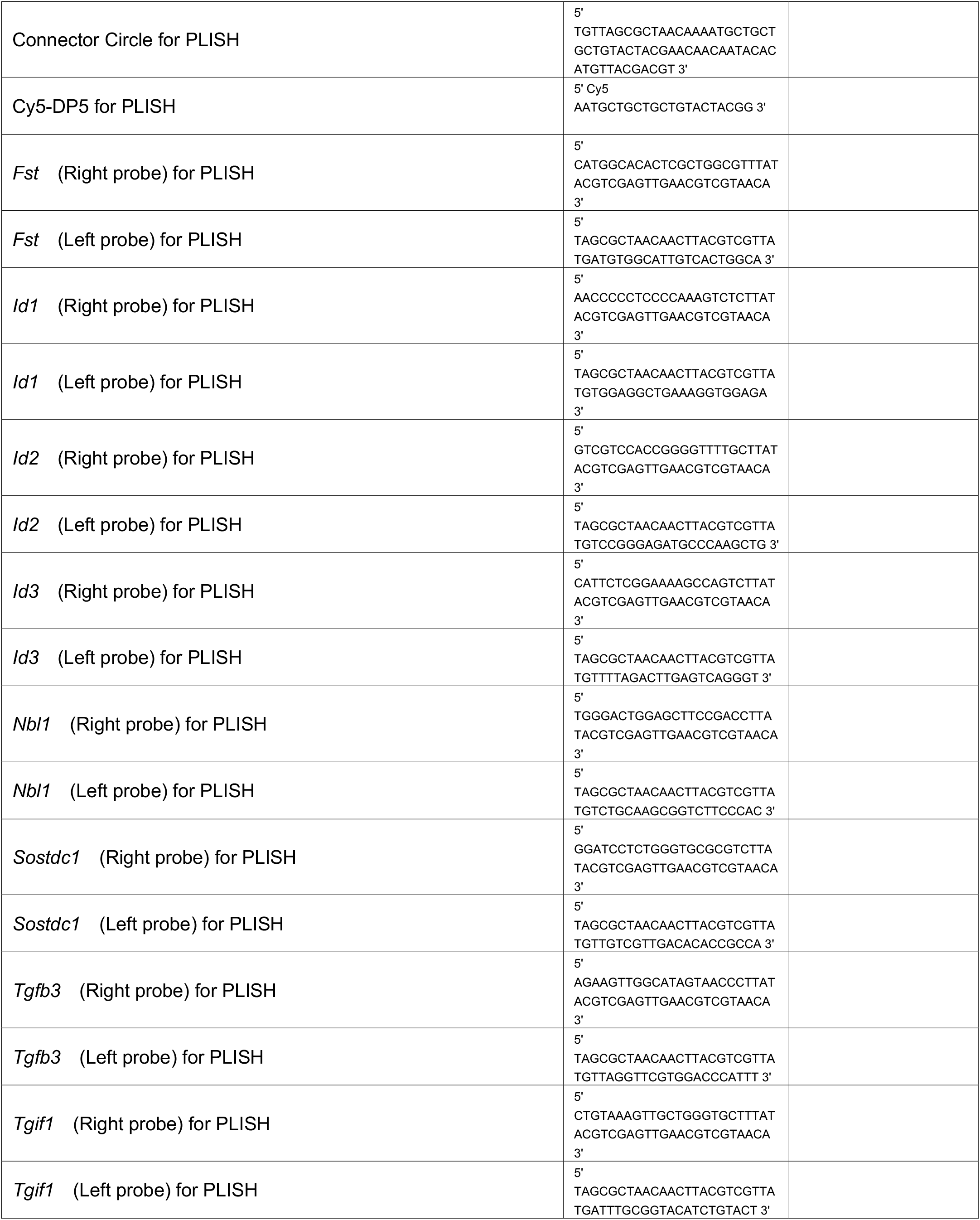
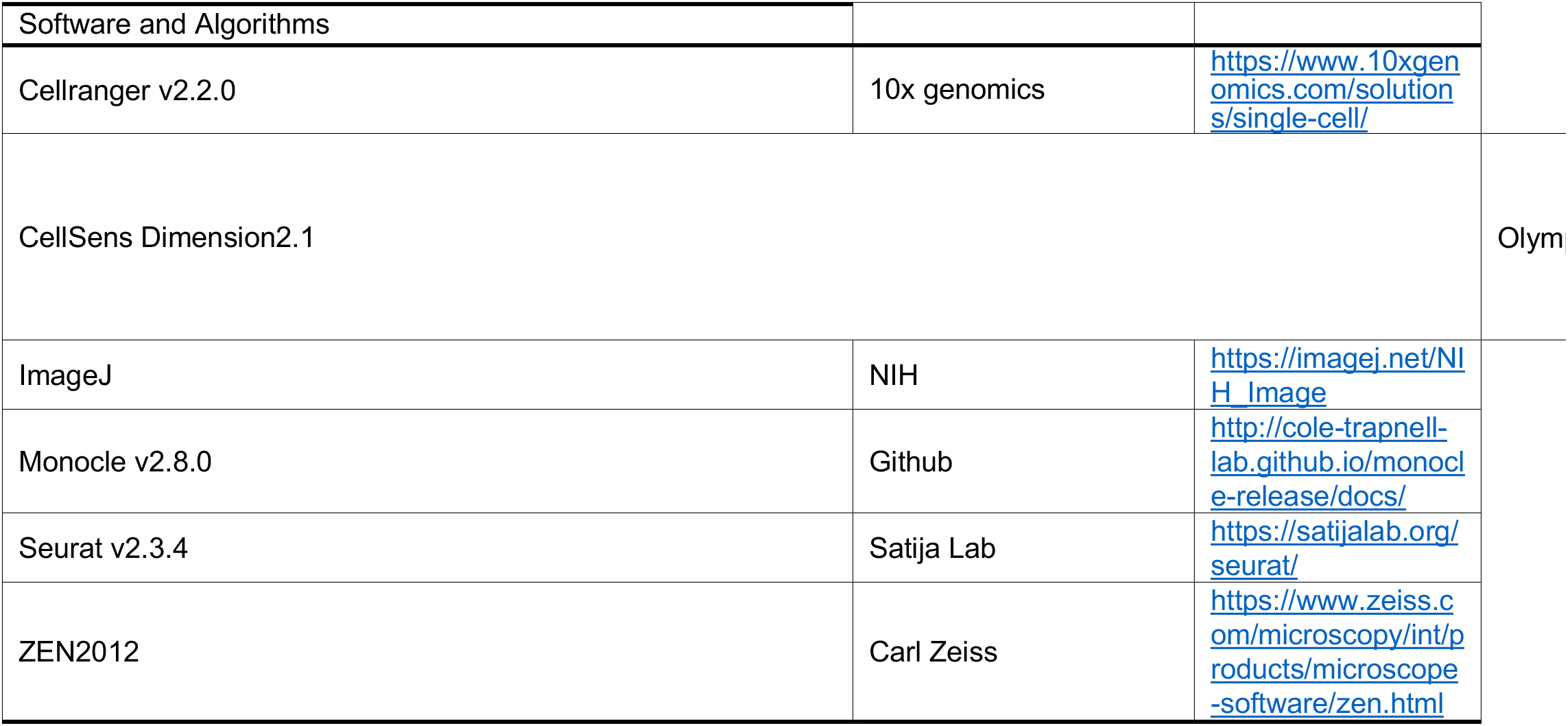

### LEAD CONTACT AND MATERIALS AVAILABILITY

Further information and requests for resources and reagents should be directed to and will be fulfilled by the Lead Contact, Dr. Mitsuru Morimoto. This study did not generate new unique reagents.

### EXPERIMENTAL MODEL

#### MICE

The Institutional Animal Care and Use Committee of RIKEN Kobe Branch approved all the experimental procedures using animal in accordance with the ethics guidelines of the institute. All mouse strains were maintained in the RIKEN BDR animal facility in specific pathogen free (SPF) conditions.

See **KEY RESOURCES TABLE** for information of each mice line. To minimalize tissue deformation, in all experiments, embryos were fixed in 4% paraformaldehyde/phosphate buffered saline (PBS) overnight at 4 °C or 3 hours at room temperature, and then tracheas were dissected.

#### Generation of *Rosa^CAG-LSL-3xHA-Id2-IRES-H2B-EGFP/+^* mice line

To generate a knock-in mouse expressing 3xHA-Id2 and H2B-EGFP that was conditionally incorporated into the ROSA26 locus, Gt(ROAS)26Sor^tm1CAG-LSL-3xHA-Id2-IRES-H2B-EGFP^ (Accession No. CDB0076E: http://www2.clst.riken.jp/arg/mutant%20mice%20list.html) mouse (Figure S3E), was established with CRISPR/Cas9 genome editing technology in zygotes as described previously(Abe, et al., 2020). In brief, the ROSA26 donor vector was constructed using Gateway technology (Thermo Fisher Scientific). The Gateway destination vector, named pR26-CAG-STOP-HR-DEST, was modified from pBigT(Srinivas, et al., 2001), pCAGGS(Niwa, et al., 1991), R26-H2B-EGFP HR donor vector(Abe, et al., 2020). pENTR2B-3xHA-Id2-IRES-H2B-EGFP was recloned into the destination vector using LR clonase of the Gateway technology in order to generate the donor vector. The donor vector was injected into C57BL/6 zygotes and the knock-in F0 mice were identified by PCR(Abe, et al., 2020). Genotype was determined by genetic PCR with combination of following primers; F1: 5’-AGCCGTTCACCTCGGGTA-3’, R1: 5’-GATACCGTCGACGCCACAGAG-3’ (440bp).

## METHOD DETAILS

### BrdU-incorporation assay

To label almost all of proliferating cells, BrdU (0.1 mg ml^−1^, Sigma-Aldrich, B5002) were cumulatively injected into pregnant mice four times every 2 h before sacrifice. To measure the proliferating ratio, BrdU^+^ cells were manually counted in the ventral epithelium between the 1st and 12th cartilage region, based on the sections immunostained for BrdU. Before E13.5, when the cartilage did not appear yet, the cells were counted in the entire region of the trachea in sagittal sections.

### Cell cycle analysis

For the assessment of cell cycle status in epithelial cells in developing trachea at E14.5 and E16.5, Rosa^FucciG1/+^ mice (Abe, et al., 2013) were mated with Shh^Cre^, Rosa^H2B-EGFP/+^. The developing trachea dissected from Shh^Cre^, Rosa^H2B-EGFP/FucciG1^ were digested by 0.25% Trypsin (Thermo Fisher Scientific, 25200056), 0.5mg/ml DNase I (Sigma-Aldrich, DN25), and 1x RBC lysis buffer (BioLegend, BL420301). The single cell suspension was obtained after passing cells through a 40 mm cell strainer. Then, the single cell suspension was sorted for selecting GFP+ epithelial cells using BD FACS Aria II appliance. Based on mCherry intensity, GFP+ epithelial cells were classified into three population such as mCherry^negatlve^, mCherry^moderate^, and mCherry^high^ cells. We defined that Cherry^moderate^ and mCherry^high^ cells are in G1 and G0 phases respectively in accordance with the previous paper (Abe, et al., 2013).

### Cell dissociation and FACS

To collect Krt17^+^ and Krt17^-^ cells using Krt17^EGFP^ mice at E18.5, single-cell suspensions were made through the incubation of trachea with Trypsin-EDTA (0.25%) (Thermo Fisher Scientific, 25200056) at 37°C for 60 min, followed by gentle pipetting and passage through a 40 mm cell strainer. For FACS with EpCAM-APC (Invitrogen, 17-5791-80), cells were diluted to less than 1 ×10^6^ cells/mL in PBS with 3% FBS and incubated in 1:100 EpCAM-APC (Invitrogen, 17-5791-80) or 1:100 IgG-APC isotype control (BioLegend, 400511) on ice for 30 min, followed by PBS wash. Sorting was performed on BD FACS Aria II and data analyzed with FACS Diva (BD Biosciences). Cells were collected in MTEC/Plus and cultured immediately or frozen for RNA extraction.

To prepare single-cell suspensions from adult trachea, the epithelial sheet was peeled off from mesenchymal tissue in trachea with a tungsten needle after the incubation with 400U/ml collagenase type I (Worthington Biochemical Corporation, CLS1) at 37°C for 60 min or 16U/ml Dispase (Corning) at 37°C for 40 min. Single-cell suspensions were made through the incubation with Trypsin-EDTA (0.25%) at 37°C for 60 min, followed by gentle pipetting, passage through a 40 mm cell strainer, and incubation in 1x RBC lysis at RT for 3 min. For FACS with EpCAM-APC, cells were diluted to less than 1 ×10^6^ cells/mL in PBS with 3% FBS and incubated in 1:100 EpCAM-APC or 1:100 IgG-APC isotype control on ice for 30 min, followed by PBS wash. Sorting was performed on BD FACS Aria II and data analyzed with FACS Diva. Cells were collected in MTEC/Plus and cultured immediately or frozen for RNA extraction.

Single-cell suspensions at E18.5 and 2M for tracheosphere culture were made through the incubation of trachea with Trypsin-EDTA (0.25%) (Thermo Fisher Scientific, 25200056) at 37°C for 30-60 min, followed by gentle pipetting and passage through a 40 mm cell strainer. To remove the mesenchymal cells, all the cells were incubated at 37°C for 2 hours in DMEM/Ham’s F-12 (Nacalai, 11582-05) with 5% FBS. Floating cells were collected and used as epithelial cells for the further tracheosphere culture. The day when culture experiment started was defined as day0. 500μl MTEC/Plus was added to each well and changed at Day1, 3. 10μM Y-27632 (LC laboratories, Y-5301) was supplemented at Day0 and Day1 only. The number of spheres with the diameter over 50μm per well was counted on Day4 using an inverted microscope.

To isolate basal cells at 2M for tracheosphere or two-dimensional culture, the epithelial sheet was peeled off from mesenchymal tissue in trachea with a tungsten needle after the incubation with 16U/ml Dispase (Corning) at 37°C for 40 min. Single-cell suspensions were made through the incubation with Trypsin-EDTA (0.25%) at 37°C for 20 min, followed by gentle pipetting, passage through a 40 mm cell strainer. For FACS, cells were diluted to less than 1 ×10^6^ cells/mL in PBS with 3% FBS and incubated in 1:100 EpCAM-APC and 1:200 anti-p75 NGF receptor antibody or 1:100 anti-BSI B4-FITC antibody on ice for 30 min, followed by PBS wash. In the case of anti-p75 NGF receptor antibody staining, incubation with 1:500 Alexa Fluor 488 Donkey anti-rabbit on ice for 30 min was followed. Sorting was performed on BD FACS Aria II and data analyzed with FACS Diva. Cells were collected in MTEC/Plus and cultured immediately. For tracheosphere culture, 2×10^3^ basal cells were co-cultured with 4×10^4^ fibroblast cells in 25ul Growth-factor reduced Matrigel with 75ul MTEC/Plus (total 100ul/well). 500μl MTEC/Plus was added to each well and changed at Day1, 3. 10μM Y-27632 (LC laboratories, Y-5301) was supplemented at Day0 and Day1 only. The number of spheres with the diameter over 50μm per well was counted on Day4 using an inverted microscope. For two-dimensional culture, 1.5×10^4^ basal cells in 300ul MTEC/Plus were seeded onto 24-well 0.4-μm Transwell insert (Corning, #3470) coated with collagen type I (Corning, 354236). 0.5mL MTEC/Plus was added to the lower chamber. After 2-day culture, MTEC/Plus with Tgfb inhibitor (SB431542, 10μM) (Sigma-Aldrich, 616461) or Tgfb1/2/3 ligands (10ng/ml, each) (R&D systems, 240-B/302-B2/243-B3) was placed in the upper and lower chambers. After 1-day culture, the cells on the Transwell insert were soaked in 400μl TRIzol™ Reagent (Invitrogen, 15596018) for Quantitative RT-PCR.

### Ex-vivo trachea culture experiment

The developing tracheas dissected from E12.5 embryos were transferred onto the Whatman Nuclepore™ track-etched polycarbonate membrane (Whatman, 110614) and cultured at an air-liquid interface with DMEM/Ham’s F-12 medium (Nacalai, 11582-05) supplemented with penicillin/streptomycin (Nacalai, 09367-34) and 5% FBS. The medium was changed every day in all experiments. The day when the ex-vivo trachea culture experiment started was defined as day0. 10μM Y-27632 (LC laboratories, Y-5301) was supplemented at Day0 and Day1 in all experiments. To assess the effects of cell cycle status on the proliferation and differentiation processes in the tracheal epithelium, E12.5 trachea was cultured in the medium supplemented with or without PD00332991 (300nM, 600nM, or 900nM) (Abcam, 09367-34). At Day2, the cultured trachea was transferred into 4% PFA for 30 min at 37 °C for fixation followed by PBS wash and the overnight incubation in 30% sucrose at 4°C, and then embedded in OCT compound. To assess the effects of Tgfb signaling on the proliferation and differentiation processes in the tracheal epithelium, E12.5 trachea was cultured in the various conditions such as Tgfb inhibitor treatment (SB431542, 2μM) (Sigma-Aldrich, 616461) and the combination treatment of Tgfb1/2/3 ligands (10ng/ml, each) (R&D systems, 240-B/302-B2/243-B3). At Day2, samples were collected and embedded in OCT compound in the same way described above.

### Microscopy and imaging

Tissue section immunofluorescence staining was imaged with LSM 710 confocal microscopy (Carl Zeiss). Cells were counted based on nuclear staining with DAPI (Nacalai, 11034-56) and specific cell markers of the respective cell types. Cells were counted in the ventral region of trachea epithelium using ×63/1.4 NA Oil objective.

The tracheosphere was imaged using DP73 inverted microscopy (Olympus). Optical section images (512 x 512 to 800 x 800 pixels for the X-Y plane and 100μm for Z-axis step; 25-35 sections) was taken to estimate CFEs. Image processing and analyses were performed using ImageJ (NIH), ZEN2012 (Carl Zeiss), CellSens Dimension 2.1(Olympus), and Adobe Illustrator (Adobe).

### Immunohistochemistry

For paraffin sections, dissected tracheas were dehydrated and embedded in paraffin. For frozen sections, dissected tracheas were incubation in 30% sucrose at 4°C overnight, and then embedded in OCT compound. 6μm paraffin and 9μm frozen sections were used for immunohistochemistory experiments. The sections were treated for epitope retrieval with HistVT One (Nacalai, 06380) at 90 °C for 5 min or 105°C for 15 min, permeabilized with 0.05% Tween in PBS, blocked using M.O.M. Immunodetection Kit (Vector laboratories, VEC-BMK-2202-1) for 1 hour at room temperature, then sections were incubated with primary antibodies at 4°C for overnight. Detailed procedure and antibodies of each staining were listed below.

| **Antibody,**<br>**Dilution** | **Company,**<br>**Catalog code** | **Fixative** | **Tissue**<br>**preparation** | **Antigen retrieval** | **Secondary**<br>**antibody** |
| --- | --- | --- | --- | --- | --- |
| **GFP**<br>**(1:400)** | Thermo Fisher Scientific | 4% PFA | Frozen | 90°C, 5 min in Histo<sup>VT</sup> One | Chicken Alexa488 |
| **GFP**<br>**(1:500)** | MBL international | 4% PFA | Frozen | 90°C, 5 min in Histo<sup>VT</sup> One | Rabbit Alexa488 |
| **Scgb1a1** | Santa Cruz | 4% PFA | Paraffin | 105°C, 15 min in Histo<sup>VT</sup> One | Goat Alexa594 |
| **(1:500)** |  |  |  |  |  |
| **Foxj1**<br>**(1:200)** | eBioscience | 4% PFA | Paraffin | 105°C, 15 min in Histo<sup>VT</sup> One | Mouse Alexa594 |
| **Ki67**<br>**(1:200)** | BD Biosciences | 4% PFA | Paraffin | 105°C, 15 min in Histo<sup>VT</sup> One | Mouse Alexa594 |
| **Krt17**<br>**(1:400)** | Santa Cruz | 4% PFA | Paraffin | 105°C, 15 min in Histo<sup>VT</sup> One | Mouse Alexa488 |
| **Krt17**<br>**(1:1000)** | Abcam | 4% PFA | Paraffin | 105°C, 15 min in Histo<sup>VT</sup> One | Rabbit Alexa488 |
| **Krt5**<br>**(1:1000)** | Abcam | 4% PFA | Paraffin | 105°C, 15 min in Histo<sup>VT</sup> One | Rabbit Alexa488 |
| **pSmad2/3**<br>**(1:200)** | Cell Signaling | 4% PFA | Frozen | 90°C, 5 min in Histo<sup>VT</sup> One | Rabbit Alexa488 |
| **Scgb3a2**<br>**(1:1000)** | Dr. Shioko Kimura's lab | 4% PFA | Paraffin | 105°C, 15 min in Histo<sup>VT</sup> One | Rabbit Alexa594 |
| **BrdU rat**<br>**(1:100).** | Abcam | 4% PFA | Paraffin | 105°C, 15 min in Histo<sup>VT</sup> One | Rat Alexa488 |
| **p63**<br>**(1:1000)** | Abcam | 4% PFA | Paraffin | 105°C, 15 min in Histo<sup>VT</sup> One | Mouse Alexa647 |

After washing the slides with 0.05% Tween in PBS for 3 times, the sections were incubated with secondary antibodies for 1 hour at room temperature. All of secondary antibody conjugated with Alexa Fluor 488/594/647 (Life Technology) were used at 1:500 dilutions. Nuclei were stained with DAPI (Nacalai, 11034-56). The sections were mounted with Fluoromount™ (Diagnostic Biosystems, K024).

For the detection of pSmad2/3, Can Get Signal^®^ immunostain Solution B (Toyobo life science, NKB-601) was used as the solution for blocking buffer and primary/secondary antibodies.

### Quantitative RT-PCR (qPCR)

Cells isolated by FACS were centrifuged and pelleted at 400*g* for 6min, then soaked in 400μl TRIzol™ Reagent (Invitrogen, 15596018). Total RNA was isolated with Directzol RNA MicroPrep (Zymoresearch, R2060) according to the manufacturer’s instructions. Reverse transcription reactions were performed with SuperScript™ III Reverse Transcriptase (Invitrogen, 18080) according to the manufacturer’s instructions. qRT-PCR was performed on 7500 Real-Time PCR instrument (Applied Biosystems) using THUNDERBIRD^®^ SYBR qPCR Mix (Toyobo life science, QPS-201). The mRNA levels of target genes were normalized to the Gapdh mRNA level. Primers used for qPCR are listed in KEY RESOURCES TABLE.

### Single cell RNA-seq for sequencing library construction

To prepare single-cell suspensions of epithelial cells in 6 time points (E12.5, E13.5, E14.5, E16.5, and E18.5), the epithelial sheet was peeled off from mesenchymal tissue in developing trachea with a tungsten needle after the incubation with 175U/ml collagenase type I (Worthington Biochemical Corporation, CLS1) at 37°C for 6 - 60 min. Single-cell suspensions were made through the incubation with Trypsin-EDTA (0.25%) (Thermo Fisher Scientific, 25200056) at 37°C for 15 min, then loaded onto Chromium Single Cell A Chips (10X Genomics, PN-1000009) for the Chromium Single Cell 3’ Library v2 (10X Genomics, PN-120233) according to the manufacturer’s recommendations (10X Genomics). Briefly, single-cell gel bead-in-emulsions (GEMs) were generated from loaded cell suspensions by a Chromium Controller instrument (10X Genomics). After performing GEM-reverse transcriptions (GEM-RTs), GEMs were harvested and the cDNAs were amplified and cleaned up with SPRIselect Reagent Kit (Beckman Coulter, B23318). Indexed sequencing libraries were constructed using Chromium Single Cell 3’ Library v2 (10X Genomics, PN-120233) for enzymatic fragmentation, end-repair, A-tailing, adaptor ligation, ligation cleanup, sample index PCR, and PCR cleanup. Libraries were sequenced on a HiSeq1500 (Illumina) to obtain a sequencing depth of around 50,000 reads per cells.

### Single cell RNA-seq Analysis

The packages listed below was used for processing raw sequencing data and downstream analysis; Cell Ranger version v2.2.0 (10X Genomics), Seurat version v2.3.4(Butler, et al., 2018), and Monocle v2.8.0(Qiu, et al., 2017a; Qiu, et al., 2017b; Trapnell, et al., 2014). First, the cells meeting any of the following criteria were omitted from further analyses for the quality control; <1,000 or >5,000 UMIs, > 7.5% of reads mapping to mitochondria genes, or EpCAM negative cells. For clustering, principal-component analysis was performed for dimension reduction. Top 15 principal components (PCs) were selected by using a permutation-based test implemented in Seurat and passed to t-Distributed Stochastic Neighbor Embedding (tSNE) for clustering visualization. To maintain a standard procedure for clustering, a value of 0.8 for the resolution was used. For the clustering analysis based on the expression of transcriptional factors, the gene list of mouse 1385 transcriptional factors derived from FANTOM5 SSTAR dataset (https://fantom.gsc.riken.jp/5/sstar/Browse_Transcription_Factors_mm9) (Lizio, et al., 2015) was used. Top 15 PCs and a value of 0.8 for the resolution were used for further clustering.

To delineate the developmental trajectories of lung progenitors in trachea, the Monocle2 algorithm (Qiu, et al., 2017a; Qiu, et al., 2017b; Trapnell, et al., 2014) was applied to the single cell dataset. Genes to be used for dimension reduction and ordering of the cells were determined by using the differentialGeneTest function in Monocle. The genes with a q-value < 0.01 were selected, and then sorted by q-value. The ordering gene set was used to compute a pseudotime graph by using the reduceDimension function (using the DDRTree method), followed by the orderCells function. To estimate the comprehensive linage map of the epithelial progenitors (Figure 1C), the unsupervised analysis was conducted without any specific cell markers. To estimate the developmental trajectories of Krt17+ and Krt17-progenitors (Figure 1H and 1I), first, the cells in S and G2-M phases were omitted. Then, the semi-supervised analysis was conducted based on Krt17 expression.

### Single molecule in-situ hybridization (PLISH)

Single molecule in situ hybridization of mRNAs called PLISH (Proximity Ligation In Situ Hybridization) was performed by following the procedures described in the past paper (Nagendran, et al., 2018). Briefly, OCT-embedded, frozen 9μm tissue sections are used to hybridize with anti-sense probe pairs that anneal at adjacent positions in a tiled manner along a target transcript. After the subsequent addition of circle and bridge oligonucleotides (circle components) harboring a specific ‘barcode’ sequence, the nicks in the junction were sealed by ligation with T4 DNA Ligase (New England BioLabs, M0202T) to create a covalently closed circle. Using the circularized probes as a template, complementary tandem repeats were generated through rolling-circle amplification (RCA) with NxGen^®^ phi29 DNA Polymerase (Lucigen, 30221). The single-stranded amplicons were detected with a Cy5-labeled oligonucleotide (Cy5-DP5) that is complementary to the specific ‘barcode’. The sets of probe pairs, circle components, and Cy5-labele oligonucleotide used to detect transcripts of the indicated genes are listed in KEY RESOURCES TABLE.

### Single molecule in-situ hybridization (RNAscope)

Single molecule in situ hybridization of mRNAs called RNAscope was performed by following the manufacturer’s instructions. Briefly, OCT-embedded, frozen 9μm tissue sections are used to hybridize with the proprietary probes for *Id2* RNA using RNAscope Multiplex Fluorescent Reagent Kit v2 (Advanced Cell Diagnostics).

### SO_2_ airway injury model

SO_2_ injury models have been previously described (Pardo-Saganta, et al., 2015; Rawlins, et al., 2007; Borthwick, et al., 2001). Briefly, 8-16 weeks old male mice were exposed to 700 ppm SO_2_ for 4 hours. Age-matched mice were used for both control and mutant mice (Shh^Cre^, Rosa^3xHA-Id2-IRES-H2B-EGFP^ mice; Id2 OE or Shh^Cre^, Tgfbr2^flox/flox^ mice; Tgfbr2 cKO). Mouse tracheas were collected 12, 18, 24, 48, 72, and 120 hours after injury. 3-4 tracheas at each time point were analyzed.

## STATISTICAL ANALYSIS

Statistical analyses were performed with Microsoft Excel for Mac. For paired comparisons, statistical significance was determined by Student’s t-test. For multiple comparisons, statistical significance was determined by Tukey’s method.

## DATA AND CORE AVAILABILITY

The scRNA-seq datasets in this paper is accessible at GSE152692. Token is itulegeoxtkzxet. (https://www.ncbi.nlm.nih.gov/geo/query/acc.cgi?acc=GSE152692)

