## Supplemental Table1 for "Airway tissue stem cells reutilize the embryonic proliferation regulator, Tgfß-Id2 axis, for tissue regeneration"

Basal markers

| Gene | average logFC | pct.1 | pct.2 | p value | Adjusted p value |
|---|---|---|---|---|---|
| Krt5 | 2.2487867 | 0.988 | 0.081 | 5.20E-299 | 8.71E-295 |
| Aqp4 | 1.2574179 | 0.797 | 0.073 | 2.11E-210 | 3.53E-206 |
| Pthlh | 2.0626199 | 0.826 | 0.094 | 4.34E-195 | 7.26E-191 |
| Adm | 2.2065094 | 0.773 | 0.081 | 5.15E-195 | 8.62E-191 |
| Col17a1 | 0.9524693 | 0.733 | 0.073 | 3.08E-176 | 5.16E-172 |
| Aqp3 | 1.803092 | 0.948 | 0.17 | 3.56E-168 | 5.96E-164 |
| Edn2 | 2.2754301 | 0.64 | 0.062 | 2.00E-159 | 3.34E-155 |
| Gm20186 | 1.2696441 | 0.68 | 0.077 | 6.01E-155 | 1.01E-150 |
| Hcar2 | 0.9493833 | 0.622 | 0.064 | 6.78E-146 | 1.14E-141 |
| Col7a1 | 0.9471133 | 0.715 | 0.095 | 3.61E-140 | 6.05E-136 |
| F3 | 2.4001233 | 1 | 0.338 | 4.69E-136 | 7.85E-132 |
| Aqp5 | 0.9072995 | 0.355 | 0.016 | 5.57E-134 | 9.32E-130 |
| Dlk2 | 1.0330998 | 0.709 | 0.103 | 8.51E-130 | 1.42E-125 |
| 8430408G22Rik | 1.8028229 | 0.826 | 0.166 | 6.12E-121 | 1.02E-116 |
| Ier3 | 2.5241612 | 0.93 | 0.296 | 1.36E-120 | 2.27E-116 |
| Ndufa4l2 | 1.9305848 | 0.983 | 0.353 | 5.76E-120 | 9.65E-116 |
| Serpine1 | 1.0134641 | 0.448 | 0.039 | 2.95E-114 | 4.93E-110 |
| Dcn | 1.7562441 | 0.977 | 0.319 | 5.58E-114 | 9.35E-110 |
| Tacstd2 | 1.4977308 | 0.953 | 0.345 | 1.81E-109 | 3.03E-105 |
| Hspb1 | 1.72385 | 1 | 0.477 | 2.04E-106 | 3.41E-102 |
| Krt15 | 2.161148 | 1 | 0.494 | 1.29E-105 | 2.16E-101 |
| Igfbp7 | 1.4162909 | 0.884 | 0.28 | 2.30E-105 | 3.85E-101 |
| Cbr2 | 2.232231 | 1 | 0.488 | 8.74E-104 | 1.46E-99 |
| Krt17 | 1.9812274 | 1 | 0.46 | 2.26E-103 | 3.78E-99 |
| Itgb4 | 0.9028684 | 0.826 | 0.214 | 2.85E-103 | 4.77E-99 |
| Pdpn | 1.22927 | 0.901 | 0.301 | 6.10E-103 | 1.02E-98 |
| Ckmt1 | 0.9594887 | 0.773 | 0.177 | 7.16E-99 | 1.20E-94 |
| Igfbp3 | 1.2220565 | 0.68 | 0.131 | 9.78E-97 | 1.64E-92 |
| Smoc2 | 0.9063398 | 0.576 | 0.087 | 5.12E-95 | 8.58E-91 |
| BC048546 | 1.6319038 | 0.872 | 0.234 | 9.22E-95 | 1.54E-90 |
| Ifitm3 | 1.7171094 | 0.988 | 0.678 | 3.74E-94 | 6.27E-90 |
| Cav1 | 1.137153 | 0.89 | 0.281 | 5.49E-94 | 9.20E-90 |
| Abi3bp | 1.4748673 | 0.884 | 0.304 | 2.47E-93 | 4.13E-89 |
| Sdc1 | 1.3532003 | 1 | 0.755 | 2.64E-92 | 4.42E-88 |
| Selenbp1 | 1.5466595 | 0.994 | 0.513 | 3.59E-92 | 6.00E-88 |
| Tmsb4x | 0.9979118 | 1 | 0.999 | 3.37E-91 | 5.65E-87 |
| Tmem176b | 1.252275 | 0.994 | 0.828 | 3.57E-91 | 5.99E-87 |
| Icam1 | 1.3522118 | 0.919 | 0.391 | 1.36E-90 | 2.28E-86 |
| Gsn | 1.0145069 | 0.86 | 0.279 | 4.52E-90 | 7.57E-86 |
| Itm2b | 1.1126936 | 1 | 0.961 | 6.36E-90 | 1.07E-85 |
| Ndrg1 | 1.0526094 | 0.855 | 0.27 | 8.73E-90 | 1.46E-85 |
| Dst | 1.101562 | 0.855 | 0.278 | 1.47E-89 | 2.47E-85 |
| Ndrg2 | 0.9480588 | 0.808 | 0.235 | 2.93E-89 | 4.91E-85 |

| | | | | | |
|---|---|---|---|---|---|
| Perp | 1.2526157 | 0.994 | 0.734 | 4.59E-89 | 7.69E-85 |
| Csrp1 | 1.3664476 | 0.971 | 0.605 | 1.38E-87 | 2.31E-83 |
| Tmem176a | 1.3602757 | 0.971 | 0.573 | 1.21E-86 | 2.02E-82 |
| Tinagl1 | 1.062208 | 0.826 | 0.274 | 1.80E-85 | 3.02E-81 |
| Krt75 | 1.104918 | 0.727 | 0.182 | 9.19E-85 | 1.54E-80 |
| Nbl1 | 1.2810416 | 0.959 | 0.455 | 4.84E-84 | 8.10E-80 |
| Cd9 | 1.0599591 | 1 | 0.928 | 5.67E-83 | 9.49E-79 |
| Cyp2f2 | 1.3150729 | 0.988 | 0.413 | 1.04E-81 | 1.75E-77 |
| Anxa3 | 1.1181121 | 0.878 | 0.339 | 1.32E-81 | 2.21E-77 |
| Actg1 | 0.8862791 | 1 | 1 | 3.34E-81 | 5.60E-77 |
| Cnn2 | 1.1672715 | 0.907 | 0.422 | 2.44E-80 | 4.09E-76 |
| Ctsl | 1.1475839 | 0.977 | 0.733 | 3.99E-79 | 6.67E-75 |
| Gsto1 | 1.4250652 | 0.983 | 0.59 | 1.01E-77 | 1.69E-73 |
| Dapl1 | 1.1088539 | 0.994 | 0.482 | 1.61E-76 | 2.70E-72 |
| Wfdc2 | 1.0693076 | 1 | 0.984 | 3.20E-76 | 5.36E-72 |
| Krt14 | 1.2222524 | 0.267 | 0.02 | 5.94E-73 | 9.94E-69 |
| Apoe | 1.0240057 | 0.436 | 0.065 | 2.43E-72 | 4.07E-68 |
| Upk1b | 0.8566736 | 0.669 | 0.174 | 1.71E-71 | 2.86E-67 |
| Phlda1 | 1.1665067 | 0.744 | 0.24 | 2.95E-69 | 4.94E-65 |
| Ifitm1 | 1.3141089 | 0.959 | 0.756 | 2.11E-68 | 3.53E-64 |
| Clic3 | 0.9080131 | 0.686 | 0.195 | 2.25E-68 | 3.76E-64 |
| 4930523C07Rik | 1.0915035 | 0.826 | 0.355 | 6.25E-68 | 1.05E-63 |
| Atp1b1 | 1.0602726 | 0.988 | 0.792 | 2.78E-67 | 4.65E-63 |
| Dusp6 | 1.1312997 | 0.878 | 0.409 | 3.83E-67 | 6.41E-63 |
| Sparc | 0.9978027 | 1 | 0.931 | 4.75E-67 | 7.95E-63 |
| Upk3bl | 1.2036584 | 0.907 | 0.448 | 2.86E-65 | 4.79E-61 |
| Tpm1 | 0.9853784 | 0.965 | 0.811 | 3.85E-65 | 6.44E-61 |
| Lmna | 1.1014387 | 0.919 | 0.549 | 1.21E-64 | 2.02E-60 |
| Fbxo32 | 1.0265254 | 0.686 | 0.215 | 2.62E-63 | 4.39E-59 |
| Gsta4 | 0.9867557 | 0.994 | 0.766 | 4.29E-63 | 7.19E-59 |
| Bsg | 0.8642545 | 1 | 0.988 | 7.00E-63 | 1.17E-58 |
| Map1lc3a | 0.9698182 | 0.977 | 0.843 | 1.15E-61 | 1.93E-57 |
| Emp2 | 0.9653105 | 0.93 | 0.629 | 3.97E-61 | 6.65E-57 |
| S100a10 | 0.9421779 | 0.977 | 0.809 | 1.56E-60 | 2.61E-56 |
| Anxa1 | 1.0257604 | 1 | 0.904 | 5.41E-60 | 9.06E-56 |
| Anxa8 | 0.8563067 | 0.977 | 0.635 | 1.00E-58 | 1.68E-54 |
| Mt1 | 0.8782491 | 0.843 | 0.358 | 8.36E-58 | 1.40E-53 |
| Tsc22d3 | 1.0815482 | 0.802 | 0.403 | 2.64E-54 | 4.43E-50 |
| Sfn | 1.0641797 | 1 | 0.878 | 1.52E-53 | 2.54E-49 |
| Cdc42ep3 | 0.9603179 | 0.942 | 0.652 | 4.81E-53 | 8.05E-49 |
| Lgals3 | 0.8689842 | 0.698 | 0.233 | 4.64E-52 | 7.76E-48 |
| Zfp36 | 1.2445906 | 0.843 | 0.485 | 1.51E-51 | 2.53E-47 |
| Sdc4 | 0.990924 | 0.895 | 0.633 | 2.89E-48 | 4.85E-44 |
| Ptrf | 0.866882 | 0.779 | 0.416 | 1.62E-47 | 2.72E-43 |
| Gprc5a | 0.956456 | 0.767 | 0.368 | 1.63E-47 | 2.73E-43 |

| | | | | | |
|---|---|---|---|---|---|
| Junb | 1.0042846 | 0.994 | 0.907 | 7.14E-47 | 1.20E-42 |
| Tnfrsf12a | 0.9290048 | 0.797 | 0.43 | 2.16E-45 | 3.62E-41 |
| Hbegf | 1.3335644 | 0.599 | 0.213 | 2.43E-45 | 4.07E-41 |
| Nupr1 | 1.0428774 | 0.587 | 0.174 | 2.12E-42 | 3.56E-38 |
| Errfi1 | 1.1759175 | 0.75 | 0.428 | 4.74E-41 | 7.94E-37 |
| Mt2 | 0.8923516 | 0.593 | 0.203 | 7.80E-41 | 1.31E-36 |
| Atf3 | 1.3687235 | 0.919 | 0.694 | 3.17E-35 | 5.30E-31 |
| Cyr61 | 1.1048126 | 0.919 | 0.768 | 4.49E-31 | 7.52E-27 |
| Socs3 | 1.1784385 | 0.64 | 0.322 | 6.12E-29 | 1.02E-24 |
| Klf4 | 0.8945324 | 0.68 | 0.444 | 2.78E-23 | 4.66E-19 |
| Nfkbia | 1.2241469 | 0.808 | 0.663 | 2.16E-22 | 3.61E-18 |
| Dusp1 | 0.942976 | 0.901 | 0.796 | 2.39E-22 | 4.00E-18 |

Club cell markers

| Gene | average logFC | pct.1 | pct.2 | p value | Adjusted p value |
|---|---|---|---|---|---|
| Scgb1a1 | 3.9746903 | 0.921 | 0.026 | 0.00E+00 | 0.00E+00 |
| AA467197 | 1.7767814 | 0.73 | 0.007 | 0.00E+00 | 0.00E+00 |
| Gp2 | 1.2070067 | 0.584 | 0.003 | 0.00E+00 | 0.00E+00 |
| Cp | 1.3469619 | 0.753 | 0.017 | 4.34E-300 | 7.26E-296 |
| Sftpd | 1.2411177 | 0.663 | 0.012 | 1.25E-294 | 2.09E-290 |
| Tff2 | 2.2753276 | 0.528 | 0.006 | 8.49E-278 | 1.42E-273 |
| Chad | 2.0029732 | 1 | 0.059 | 4.06E-242 | 6.80E-238 |
| Ltf | 1.3375851 | 0.629 | 0.016 | 2.58E-235 | 4.33E-231 |
| Creb3l1 | 1.1550509 | 0.944 | 0.056 | 3.41E-216 | 5.72E-212 |
| Muc16 | 1.2464643 | 0.73 | 0.029 | 3.80E-216 | 6.36E-212 |
| Muc5b | 3.593379 | 0.944 | 0.075 | 1.09E-186 | 1.83E-182 |
| Chil1 | 0.8759478 | 0.393 | 0.006 | 1.52E-182 | 2.55E-178 |
| Lcn2 | 1.3128433 | 0.404 | 0.007 | 1.01E-176 | 1.69E-172 |
| Cd36 | 0.7854814 | 0.652 | 0.03 | 7.76E-173 | 1.30E-168 |
| Scgb3a1 | 5.4214119 | 1 | 0.11 | 5.28E-163 | 8.85E-159 |
| Lyz2 | 0.8551394 | 0.292 | 0.003 | 2.90E-153 | 4.85E-149 |
| Agr2 | 3.7431954 | 0.978 | 0.116 | 1.52E-146 | 2.54E-142 |
| Ly6a | 0.8344162 | 0.73 | 0.051 | 4.91E-138 | 8.22E-134 |
| Sftpb | 1.6077694 | 0.382 | 0.011 | 4.03E-132 | 6.74E-128 |
| Ptges | 0.8974749 | 0.955 | 0.11 | 1.59E-124 | 2.66E-120 |
| Trf | 2.5148272 | 0.989 | 0.148 | 5.39E-122 | 9.03E-118 |
| Pglyrp1 | 3.2531157 | 1 | 0.174 | 1.35E-118 | 2.26E-114 |
| 5330417C22Rik | 0.8582244 | 0.876 | 0.102 | 7.42E-117 | 1.24E-112 |
| Nupr1 | 2.4864661 | 1 | 0.173 | 1.33E-111 | 2.22E-107 |
| H2-K1 | 1.0923072 | 0.933 | 0.142 | 4.54E-108 | 7.61E-104 |
| Klk13 | 0.9421191 | 0.551 | 0.039 | 3.17E-107 | 5.30E-103 |
| Msln | 1.333797 | 0.876 | 0.119 | 3.70E-102 | 6.20E-98 |
| Lrrc26 | 1.0439942 | 0.966 | 0.156 | 8.05E-95 | 1.35E-90 |
| AU021092 | 1.8886746 | 1 | 0.231 | 1.63E-94 | 2.73E-90 |
| Reg3g | 4.8567038 | 1 | 0.249 | 5.38E-93 | 9.01E-89 |
| Fkbp11 | 1.177135 | 0.685 | 0.087 | 5.13E-87 | 8.59E-83 |
| Wfdc1 | 0.7319007 | 0.573 | 0.053 | 3.75E-86 | 6.29E-82 |
| Muc1 | 0.8432451 | 0.809 | 0.128 | 1.10E-80 | 1.85E-76 |
| Hp | 1.5747356 | 0.618 | 0.07 | 8.81E-79 | 1.48E-74 |
| Tspan1 | 1.5336298 | 0.989 | 0.276 | 2.76E-76 | 4.63E-72 |
| Klk10 | 2.2106622 | 0.753 | 0.125 | 8.24E-76 | 1.38E-71 |
| Isg20 | 1.271994 | 0.944 | 0.24 | 8.25E-76 | 1.38E-71 |
| Ly6e | 2.349996 | 1 | 0.36 | 2.55E-75 | 4.27E-71 |
| Sec11c | 1.2963959 | 0.978 | 0.298 | 4.17E-73 | 6.98E-69 |
| Appl2 | 0.8333372 | 0.899 | 0.19 | 5.17E-71 | 8.67E-67 |
| Shisa5 | 1.0033202 | 0.978 | 0.285 | 1.14E-65 | 1.91E-61 |

| | | | | | |
|---|---|---|---|---|---|
| Aldh1a1 | 1.2582266 | 0.978 | 0.292 | 6.34E-65 | 1.06E-60 |
| Cldn10 | 1.398248 | 1 | 0.335 | 3.74E-62 | 6.27E-58 |
| Fgfbp1 | 1.1564416 | 0.933 | 0.264 | 6.35E-62 | 1.06E-57 |
| Scgb3a2 | 3.6529489 | 1 | 0.578 | 1.05E-61 | 1.76E-57 |
| Dcxr | 1.3834215 | 1 | 0.409 | 2.27E-61 | 3.79E-57 |
| Nucb2 | 0.962384 | 0.876 | 0.236 | 9.05E-61 | 1.52E-56 |
| Mgst2 | 0.7887568 | 0.73 | 0.139 | 2.18E-59 | 3.64E-55 |
| Oit1 | 1.081398 | 0.978 | 0.305 | 1.29E-58 | 2.17E-54 |
| Lypd2 | 1.9263188 | 1 | 0.651 | 5.69E-57 | 9.53E-53 |
| Qsox1 | 1.2784431 | 0.966 | 0.368 | 6.40E-56 | 1.07E-51 |
| Tmed3 | 1.562748 | 0.989 | 0.51 | 1.36E-55 | 2.28E-51 |
| Selenbp1 | 2.0906729 | 1 | 0.524 | 1.92E-55 | 3.21E-51 |
| Ssr4 | 1.7691126 | 0.989 | 0.768 | 6.48E-55 | 1.08E-50 |
| Gfpt1 | 0.9798076 | 0.809 | 0.21 | 8.38E-55 | 1.40E-50 |
| Gsto1 | 2.0419676 | 1 | 0.599 | 1.69E-54 | 2.84E-50 |
| Wfdc2 | 1.6248566 | 1 | 0.985 | 3.44E-54 | 5.76E-50 |
| H2-D1 | 1.1121006 | 0.921 | 0.34 | 1.96E-52 | 3.28E-48 |
| Tcn2 | 0.7986825 | 0.831 | 0.214 | 9.05E-52 | 1.51E-47 |
| 8430408G22Rik | 1.1721202 | 0.809 | 0.182 | 5.69E-51 | 9.53E-47 |
| Cbr2 | 1.8063753 | 1 | 0.501 | 5.58E-49 | 9.34E-45 |
| F3 | 1.1948289 | 0.989 | 0.355 | 2.88E-48 | 4.83E-44 |
| Bsg | 1.2195909 | 1 | 0.988 | 5.88E-47 | 9.85E-43 |
| Ndufa4l2 | 1.1555167 | 1 | 0.367 | 1.03E-46 | 1.72E-42 |
| Cited1 | 1.251802 | 0.831 | 0.276 | 1.12E-46 | 1.87E-42 |
| Tmbim4 | 0.9614934 | 0.944 | 0.456 | 2.25E-46 | 3.76E-42 |
| Aldh1a7 | 0.9076989 | 0.978 | 0.386 | 7.63E-46 | 1.28E-41 |
| Mien1 | 1.1502708 | 0.989 | 0.641 | 8.15E-45 | 1.36E-40 |
| Spint2 | 0.7736045 | 1 | 0.979 | 1.04E-44 | 1.74E-40 |
| Ddost | 0.9564703 | 1 | 0.79 | 1.16E-44 | 1.95E-40 |
| Mgst1 | 1.507352 | 1 | 0.868 | 1.49E-44 | 2.50E-40 |
| Gale | 0.7699339 | 0.854 | 0.276 | 3.14E-43 | 5.25E-39 |
| Plpp3 | 0.790726 | 0.865 | 0.266 | 3.73E-42 | 6.24E-38 |
| Edem2 | 0.7980422 | 0.978 | 0.439 | 1.27E-41 | 2.12E-37 |
| P4hb | 1.0308796 | 0.978 | 0.815 | 1.72E-41 | 2.89E-37 |
| Sec61b | 0.85747 | 1 | 0.934 | 4.19E-41 | 7.01E-37 |
| Ppib | 0.9622653 | 1 | 0.955 | 4.03E-40 | 6.75E-36 |
| Upk3a | 0.7599255 | 0.292 | 0.03 | 2.89E-39 | 4.84E-35 |
| Serp1 | 0.9953552 | 0.955 | 0.711 | 6.51E-39 | 1.09E-34 |
| Cyp2f2 | 1.2513369 | 1 | 0.426 | 8.09E-39 | 1.36E-34 |
| Ssr2 | 0.8915395 | 1 | 0.921 | 8.15E-39 | 1.36E-34 |
| Smim14 | 0.941058 | 0.966 | 0.637 | 1.47E-38 | 2.46E-34 |
| Rabac1 | 1.0500381 | 0.944 | 0.701 | 6.03E-38 | 1.01E-33 |
| Ankrd37 | 0.9070236 | 0.775 | 0.248 | 2.55E-37 | 4.27E-33 |

| | | | | | |
|---|---|---|---|---|---|
| Tspan13 | 0.9637662 | 0.966 | 0.689 | 6.20E-37 | 1.04E-32 |
| Krtcap2 | 0.8125518 | 0.978 | 0.812 | 2.90E-36 | 4.86E-32 |
| Fkbp2 | 0.7622263 | 0.955 | 0.581 | 1.52E-35 | 2.55E-31 |
| Dad1 | 0.7334801 | 0.989 | 0.878 | 4.25E-35 | 7.11E-31 |
| Tst | 0.812851 | 0.966 | 0.488 | 4.19E-34 | 7.01E-30 |
| Ctsh | 0.9701358 | 0.933 | 0.5 | 3.22E-33 | 5.40E-29 |
| Krt18 | 0.9666311 | 1 | 0.933 | 7.03E-33 | 1.18E-28 |
| BC048546 | 0.9453012 | 0.831 | 0.25 | 1.10E-32 | 1.83E-28 |
| Edn2 | 1.0852631 | 0.438 | 0.082 | 1.02E-31 | 1.71E-27 |
| Calr | 0.7244209 | 0.989 | 0.957 | 3.01E-31 | 5.03E-27 |
| Atp1b1 | 0.8109048 | 1 | 0.796 | 8.24E-31 | 1.38E-26 |
| Rrbp1 | 0.724226 | 0.944 | 0.724 | 1.16E-29 | 1.94E-25 |
| Gsta4 | 0.8416858 | 1 | 0.771 | 1.07E-28 | 1.79E-24 |
| Manf | 0.8308531 | 0.966 | 0.817 | 2.91E-26 | 4.88E-22 |
| Meg3 | 0.9644072 | 0.955 | 0.877 | 5.99E-21 | 1.00E-16 |
| Krt15 | 0.8865383 | 0.921 | 0.508 | 2.02E-18 | 3.39E-14 |

Ciliated cell markers

| Gene | average logFC | pct.1 | pct.2 | p value | Adjusted p value |
| --- | --- | --- | --- | --- | --- |
| Dynlrb2 | 3.1621075 | 0.966 | 0.012 | 0.00E+00 | 0.00E+00 |
| Ccdc153 | 3.1085458 | 0.672 | 0.006 | 0.00E+00 | 0.00E+00 |
| Hdc | 2.8296287 | 0.741 | 0.005 | 0.00E+00 | 0.00E+00 |
| Rsph1 | 2.6787606 | 0.931 | 0.017 | 0.00E+00 | 0.00E+00 |
| 1110017D15Rik | 2.6633311 | 0.931 | 0.008 | 0.00E+00 | 0.00E+00 |
| Tmem212 | 2.5030619 | 0.672 | 0.003 | 0.00E+00 | 0.00E+00 |
| Meig1 | 2.4727272 | 0.931 | 0.01 | 0.00E+00 | 0.00E+00 |
| Fam183b | 2.4041489 | 0.845 | 0.008 | 0.00E+00 | 0.00E+00 |
| Pifo | 2.3394968 | 0.966 | 0.019 | 0.00E+00 | 0.00E+00 |
| Efcab10 | 2.2420142 | 0.81 | 0.007 | 0.00E+00 | 0.00E+00 |
| 3300002A11Rik | 2.1939733 | 0.603 | 0.001 | 0.00E+00 | 0.00E+00 |
| Lrrc23 | 2.1384576 | 0.828 | 0.005 | 0.00E+00 | 0.00E+00 |
| Sec14l3 | 2.1045804 | 0.793 | 0.008 | 0.00E+00 | 0.00E+00 |
| Tekt1 | 2.0839345 | 0.931 | 0.018 | 0.00E+00 | 0.00E+00 |
| Tctex1d4 | 2.0692747 | 0.638 | 0.004 | 0.00E+00 | 0.00E+00 |
| Morn5 | 1.9983484 | 0.828 | 0.005 | 0.00E+00 | 0.00E+00 |
| 1700001C02Rik | 1.9786403 | 0.776 | 0.002 | 0.00E+00 | 0.00E+00 |
| Dnali1 | 1.9531384 | 0.828 | 0.011 | 0.00E+00 | 0.00E+00 |
| Sntn | 1.8775846 | 0.603 | 0.001 | 0.00E+00 | 0.00E+00 |
| 2610028H24Rik | 1.8350875 | 0.81 | 0.003 | 0.00E+00 | 0.00E+00 |
| Ccdc113 | 1.8098155 | 0.862 | 0.008 | 0.00E+00 | 0.00E+00 |
| Tekt4 | 1.7467251 | 0.862 | 0.002 | 0.00E+00 | 0.00E+00 |
| Fam92b | 1.7207794 | 0.707 | 0.002 | 0.00E+00 | 0.00E+00 |
| Vpreb3 | 1.6973279 | 0.621 | 0.001 | 0.00E+00 | 0.00E+00 |
| BC051019 | 1.6760487 | 0.828 | 0.003 | 0.00E+00 | 0.00E+00 |
| Odf3b | 1.6759937 | 0.759 | 0.003 | 0.00E+00 | 0.00E+00 |
| Cfap206 | 1.6431138 | 0.931 | 0.014 | 0.00E+00 | 0.00E+00 |
| Zmynd10 | 1.6390105 | 0.897 | 0.005 | 0.00E+00 | 0.00E+00 |
| Morn3 | 1.6272945 | 0.759 | 0.001 | 0.00E+00 | 0.00E+00 |
| Dnah12 | 1.6133061 | 0.897 | 0.013 | 0.00E+00 | 0.00E+00 |
| Gm867 | 1.5933885 | 0.603 | 0.001 | 0.00E+00 | 0.00E+00 |
| 1700012B09Rik | 1.5743669 | 0.672 | 0.001 | 0.00E+00 | 0.00E+00 |
| Stmnd1 | 1.5621567 | 0.603 | 2.00E-03 | 0.00E+00 | 0.00E+00 |
| 2410004P03Rik | 1.5417049 | 0.793 | 6.00E-03 | 0.00E+00 | 0.00E+00 |
| 4833427G06Rik | 1.5054138 | 0.845 | 2.00E-03 | 0.00E+00 | 0.00E+00 |
| Ccdc67 | 1.4719 | 0.69 | 8.00E-03 | 0.00E+00 | 0.00E+00 |
| Dnah5 | 1.4545704 | 0.69 | 1.00E-03 | 0.00E+00 | 0.00E+00 |
| 1700001L19Rik | 1.4287999 | 0.776 | 3.00E-03 | 0.00E+00 | 0.00E+00 |
| Ccdc78 | 1.5015812 | 0.621 | 6.00E-03 | 2.97E-293 | 4.98E-289 |

| | | | | | |
|---|---|---|---|---|---|
| Prr29 | 1.5591636 | 0.672 | 1.00E-02 | 8.82E-270 | 1.48E-265 |
| Foxj1 | 2.2826697 | 0.983 | 3.10E-02 | 2.04E-268 | 3.42E-264 |
| 1700016K19Rik | 2.6127726 | 0.983 | 3.20E-02 | 4.32E-265 | 7.23E-261 |
| Plet1 | 1.88337 | 0.793 | 1.70E-02 | 1.27E-264 | 2.13E-260 |
| Lrrc48 | 1.6499456 | 0.897 | 2.80E-02 | 1.71E-246 | 2.86E-242 |
| Cfap53 | 1.4434423 | 0.707 | 1.50E-02 | 1.24E-237 | 2.07E-233 |
| Myl4 | 1.8077844 | 0.603 | 1.00E-02 | 9.32E-233 | 1.56E-228 |
| Agr3 | 1.638465 | 0.776 | 2.10E-02 | 2.21E-228 | 3.71E-224 |
| Riiad1 | 2.6433147 | 0.914 | 3.40E-02 | 4.61E-226 | 7.72E-222 |
| Cfap126 | 2.7442066 | 0.983 | 4.20E-02 | 5.99E-222 | 1.00E-217 |
| Nme5 | 2.0441173 | 0.966 | 4.10E-02 | 5.51E-218 | 9.23E-214 |
| Tm4sf1 | 1.9718646 | 0.81 | 2.70E-02 | 1.70E-208 | 2.85E-204 |
| Smim5 | 1.5891519 | 0.81 | 3.30E-02 | 1.97E-183 | 3.30E-179 |
| Dmkn | 2.313567 | 0.759 | 2.80E-02 | 1.23E-182 | 2.06E-178 |
| 1700026L06Rik | 2.0299222 | 0.759 | 2.80E-02 | 1.17E-180 | 1.96E-176 |
| AU040972 | 3.5007573 | 0.672 | 2.10E-02 | 2.32E-177 | 3.89E-173 |
| Ccdc17 | 1.7047179 | 0.724 | 2.60E-02 | 1.57E-176 | 2.64E-172 |
| 1700007K13Rik | 2.2926432 | 0.914 | 6.10E-02 | 2.79E-146 | 4.67E-142 |
| Cfap45 | 1.785319 | 0.966 | 8.00E-02 | 3.66E-133 | 6.13E-129 |
| Cetn4 | 1.9801996 | 0.879 | 6.60E-02 | 3.21E-127 | 5.38E-123 |
| Erich2 | 2.1957679 | 0.879 | 6.90E-02 | 1.05E-123 | 1.76E-119 |
| 1700088E04Rik | 1.748617 | 0.948 | 8.40E-02 | 1.84E-122 | 3.09E-118 |
| Ccdc189 | 1.9228339 | 0.862 | 7.30E-02 | 3.29E-112 | 5.51E-108 |
| Cmbl | 1.4553877 | 0.707 | 4.80E-02 | 2.03E-107 | 3.41E-103 |
| Fbxo36 | 1.8418576 | 0.897 | 1.06E-01 | 2.09E-90 | 3.50E-86 |
| Lrrc51 | 2.5632002 | 0.862 | 1.03E-01 | 3.76E-86 | 6.30E-82 |
| Rsph9 | 1.6304985 | 0.897 | 1.17E-01 | 5.30E-85 | 8.87E-81 |
| Spa17 | 1.9069078 | 0.879 | 1.14E-01 | 3.22E-83 | 5.40E-79 |
| Ccpg1os | 1.8245181 | 0.948 | 1.58E-01 | 8.45E-77 | 1.41E-72 |
| Spef1 | 1.5525522 | 0.862 | 1.21E-01 | 5.80E-74 | 9.71E-70 |
| Mlf1 | 2.5165639 | 1 | 2.36E-01 | 1.54E-64 | 2.58E-60 |
| Hp | 1.624784 | 0.655 | 7.40E-02 | 6.30E-63 | 1.06E-58 |
| Tppp3 | 3.4160979 | 0.983 | 2.71E-01 | 5.97E-56 | 1.00E-51 |
| Tctex1d2 | 1.5293124 | 0.948 | 2.34E-01 | 4.72E-55 | 7.90E-51 |
| Ndufaf3 | 1.4891661 | 0.966 | 2.88E-01 | 9.71E-52 | 1.63E-47 |
| Capsl | 1.8899624 | 0.966 | 2.88E-01 | 5.06E-51 | 8.47E-47 |
| Tmem107 | 1.9784482 | 0.983 | 3.47E-01 | 1.45E-49 | 2.42E-45 |
| Ccdc181 | 1.8313913 | 0.845 | 1.87E-01 | 5.40E-49 | 9.04E-45 |
| Ift22 | 1.6189464 | 1 | 4.32E-01 | 1.20E-45 | 2.01E-41 |
| 1110004E09Rik | 1.5062429 | 0.983 | 4.19E-01 | 8.73E-43 | 1.46E-38 |
| 4931406C07Rik | 1.4606598 | 0.672 | 1.24E-01 | 5.38E-42 | 9.02E-38 |

| | | | | | |
|---|---|---|---|---|---|
| Hspa4l | 1.7341689 | 0.948 | 3.79E-01 | 2.65E-41 | 4.44E-37 |
| 1110032A03Rik | 1.5492414 | 0.897 | 2.95E-01 | 1.07E-40 | 1.78E-36 |
| Cetn2 | 1.6358763 | 1 | 5.76E-01 | 3.60E-40 | 6.02E-36 |
| Dpcd | 1.7138671 | 1 | 5.67E-01 | 5.08E-39 | 8.50E-35 |
| Nudc | 1.6655432 | 1 | 8.38E-01 | 4.64E-38 | 7.76E-34 |
| Hsp90aa1 | 1.8200515 | 1 | 9.90E-01 | 1.65E-37 | 2.76E-33 |
| Elof1 | 2.3192711 | 1 | 7.66E-01 | 3.15E-37 | 5.28E-33 |
| Ruvbl2 | 1.4281199 | 0.983 | 5.42E-01 | 2.14E-36 | 3.59E-32 |
| Mt2 | 1.909324 | 0.81 | 2.12E-01 | 3.05E-36 | 5.11E-32 |
| Lgals3 | 1.8701568 | 0.845 | 2.46E-01 | 4.16E-35 | 6.96E-31 |
| Chchd6 | 1.5060701 | 0.931 | 3.99E-01 | 1.68E-33 | 2.82E-29 |
| Tubb4b | 2.8154142 | 1 | 8.05E-01 | 3.27E-33 | 5.48E-29 |
| Bphl | 1.4401011 | 0.931 | 4.12E-01 | 3.68E-33 | 6.17E-29 |
| Aldh1a1 | 1.7232663 | 0.879 | 3.00E-01 | 9.54E-33 | 1.60E-28 |
| Mt1 | 2.6933423 | 0.879 | 3.74E-01 | 4.30E-32 | 7.20E-28 |
| Chchd10 | 2.6358261 | 0.948 | 4.74E-01 | 1.09E-31 | 1.83E-27 |
| 4933434E20Rik | 2.4363004 | 0.862 | 3.82E-01 | 8.25E-29 | 1.38E-24 |
| Eln | 1.5817132 | 0.534 | 1.01E-01 | 1.66E-27 | 2.79E-23 |
| Gsn | 1.466235 | 0.741 | 3.01E-01 | 2.81E-19 | 4.71E-15 |
| Tuba1a | 2.2920797 | 0.914 | 7.02E-01 | 4.62E-19 | 7.74E-15 |

Neuroendocrine cell markers

| Gene | average logFC | pct.1 | pct.2 | p value | Adjusted p value |
|---|---|---|---|---|---|
| Ascl1 | 2.7968279 | 0.929 | 0.003 | 0.00E+00 | 0.00E+00 |
| Iapp | 2.1294049 | 0.571 | 0.001 | 0.00E+00 | 0.00E+00 |
| Chga | 1.2210294 | 0.571 | 0 | 0.00E+00 | 0.00E+00 |
| Dner | 1.1954149 | 0.929 | 0.002 | 0.00E+00 | 0.00E+00 |
| Cib3 | 1.0717909 | 0.714 | 0.001 | 0.00E+00 | 0.00E+00 |
| Crmp1 | 0.983033 | 0.786 | 0.002 | 0.00E+00 | 0.00E+00 |
| Miat | 0.9551982 | 0.714 | 0.002 | 0.00E+00 | 0.00E+00 |
| Tcerg1l | 0.8396307 | 0.786 | 0.002 | 0.00E+00 | 0.00E+00 |
| Tubb3 | 1.736089 | 0.857 | 0.004 | 1.06E-292 | 1.78E-288 |
| Scn9a | 1.1998058 | 0.714 | 0.003 | 1.29E-273 | 2.16E-269 |
| Elavl3 | 1.1424466 | 0.714 | 0.003 | 4.37E-273 | 7.31E-269 |
| Insm1 | 1.5068084 | 0.857 | 0.005 | 7.61E-264 | 1.27E-259 |
| Scg5 | 1.1927133 | 0.714 | 0.003 | 4.76E-249 | 7.97E-245 |
| Pkib | 1.2038242 | 0.786 | 0.005 | 4.24E-221 | 7.09E-217 |
| Rundc3a | 0.9229072 | 0.714 | 0.005 | 1.95E-210 | 3.27E-206 |
| Stmn3 | 1.031243 | 0.5 | 0.002 | 1.47E-191 | 2.46E-187 |
| Tmem163 | 1.4476503 | 0.929 | 0.014 | 1.54E-154 | 2.58E-150 |
| Runx1t1 | 1.4506496 | 0.714 | 0.008 | 2.48E-141 | 4.16E-137 |
| Smpd3 | 0.8657264 | 0.714 | 0.008 | 2.67E-141 | 4.48E-137 |
| Mfng | 1.0548138 | 0.786 | 0.011 | 3.81E-133 | 6.38E-129 |
| Slc38a5 | 1.2991539 | 0.643 | 0.007 | 2.76E-131 | 4.61E-127 |
| Pcsk1n | 1.1733898 | 0.571 | 0.006 | 1.66E-121 | 2.77E-117 |
| Arg1 | 1.0646891 | 0.643 | 0.009 | 6.43E-106 | 1.08E-101 |
| Lrp11 | 0.9322848 | 0.714 | 0.013 | 2.90E-97 | 4.85E-93 |
| Snca | 1.6119702 | 0.643 | 0.012 | 8.10E-90 | 1.36E-85 |
| Col8a1 | 1.3946936 | 0.714 | 0.015 | 1.45E-87 | 2.42E-83 |
| Gng3 | 0.899723 | 0.5 | 0.008 | 6.61E-77 | 1.11E-72 |
| Prox1 | 1.0805141 | 0.929 | 0.035 | 4.76E-70 | 7.97E-66 |
| Nov | 1.7525734 | 0.786 | 0.03 | 1.31E-59 | 2.20E-55 |
| Dmpk | 0.8979991 | 0.786 | 0.029 | 6.90E-59 | 1.16E-54 |
| Elavl4 | 0.9012057 | 0.571 | 0.015 | 1.97E-58 | 3.29E-54 |
| Mt3 | 0.9902706 | 0.571 | 0.015 | 4.23E-57 | 7.08E-53 |
| Rab3b | 0.8980017 | 0.571 | 0.019 | 1.56E-46 | 2.61E-42 |
| Snap25 | 0.8183301 | 0.571 | 0.021 | 4.72E-44 | 7.91E-40 |
| Hoxb5 | 1.1835147 | 0.571 | 0.024 | 9.76E-39 | 1.63E-34 |
| Ddc | 1.5275162 | 0.857 | 0.061 | 5.13E-38 | 8.59E-34 |
| Cacna2d1 | 1.4990793 | 1 | 0.089 | 1.18E-35 | 1.97E-31 |

| | | | | | |
|---|---|---|---|---|---|
| Cplx2 | 1.7623321 | 1 | 0.117 | 2.36E-29 | 3.96E-25 |
| Rtn1 | 1.0748355 | 0.286 | 0.008 | 5.02E-28 | 8.40E-24 |
| Syt7 | 1.2852149 | 0.714 | 0.063 | 9.44E-25 | 1.58E-20 |
| Pak3 | 1.0511141 | 0.714 | 0.062 | 1.06E-24 | 1.77E-20 |
| Meis2 | 2.4441104 | 0.929 | 0.139 | 1.65E-22 | 2.76E-18 |
| Tmem158 | 1.943667 | 0.929 | 0.142 | 6.53E-22 | 1.09E-17 |
| Dnajc12 | 1.5869735 | 0.929 | 0.142 | 7.94E-22 | 1.33E-17 |
| C2cd4b | 0.9708169 | 0.714 | 0.072 | 9.63E-22 | 1.61E-17 |
| Olfm1 | 0.880106 | 0.643 | 0.061 | 4.73E-21 | 7.92E-17 |
| Aplp1 | 1.1836983 | 0.714 | 0.084 | 3.12E-19 | 5.23E-15 |
| Tmod2 | 0.8324039 | 0.571 | 0.061 | 2.49E-16 | 4.17E-12 |
| Gch1 | 0.9219207 | 0.786 | 0.129 | 3.80E-15 | 6.37E-11 |
| Espn | 1.7133984 | 0.929 | 0.227 | 1.17E-14 | 1.96E-10 |
| Ncam1 | 0.7848822 | 0.643 | 0.087 | 3.75E-14 | 6.28E-10 |
| Ckb | 1.4904072 | 0.929 | 0.225 | 4.33E-13 | 7.25E-09 |
| Abcc4 | 0.9604009 | 0.357 | 0.029 | 6.01E-13 | 1.01E-08 |
| Nnat | 2.5695298 | 1 | 0.461 | 2.08E-12 | 3.49E-08 |
| Tshz2 | 0.8815623 | 0.857 | 0.22 | 9.37E-12 | 1.57E-07 |
| Cdc14b | 1.2213234 | 0.857 | 0.24 | 1.32E-11 | 2.21E-07 |
| Ly6h | 1.5963066 | 0.786 | 0.198 | 4.60E-11 | 7.70E-07 |
| Serpine2 | 0.8128358 | 0.857 | 0.212 | 5.35E-11 | 8.96E-07 |
| Bex2 | 1.3219554 | 1 | 0.339 | 6.60E-11 | 1.11E-06 |
| Uchl1 | 1.2581536 | 1 | 0.5 | 1.42E-10 | 2.38E-06 |
| Car8 | 1.0070715 | 0.643 | 0.122 | 1.87E-10 | 3.14E-06 |
| Camk2n1 | 0.891362 | 0.857 | 0.256 | 2.21E-10 | 3.71E-06 |
| Tox | 0.8994586 | 0.643 | 0.133 | 3.86E-10 | 6.47E-06 |
| Tuba1a | 1.6393454 | 1 | 0.705 | 5.75E-10 | 9.63E-06 |
| Fam46a | 1.1949942 | 0.714 | 0.176 | 1.62E-09 | 2.72E-05 |
| Hes6 | 1.5183467 | 1 | 0.77 | 1.71E-09 | 2.87E-05 |
| Gadd45g | 1.4331575 | 1 | 0.391 | 2.41E-09 | 4.04E-05 |
| Atp6v0e | 1.3715287 | 1 | 0.774 | 3.04E-09 | 5.09E-05 |
| Atp6v0b | 1.0649921 | 1 | 0.693 | 4.63E-09 | 7.76E-05 |
| Pgf | 0.853468 | 0.571 | 0.116 | 5.92E-09 | 9.91E-05 |
| Celf4 | 1.1100862 | 0.714 | 0.213 | 1.10E-08 | 1.84E-04 |
| Hoxb4 | 1.2126417 | 0.857 | 0.39 | 1.47E-07 | 2.46E-03 |
| Gdi1 | 0.8455902 | 0.929 | 0.51 | 1.62E-07 | 2.71E-03 |
| Klc1 | 1.0037993 | 0.786 | 0.342 | 2.06E-07 | 3.45E-03 |
| Selk | 0.8058518 | 1 | 0.879 | 2.20E-07 | 3.68E-03 |
| Krt7 | 1.1963507 | 0.929 | 0.896 | 2.81E-07 | 4.71E-03 |

| | | | | | |
|---|---|---|---|---|---|
| Vamp2 | 1.0500381 | 0.786 | 0.33 | 3.67E-07 | 6.14E-03 |
| Meg3 | 1.5090885 | 1 | 0.878 | 5.05E-07 | 8.46E-03 |
| Hist3h2a | 0.897404 | 1 | 0.511 | 5.93E-07 | 9.93E-03 |
| Cd24a | 1.0190144 | 1 | 0.965 | 1.40E-06 | 2.35E-02 |
| Mtpn | 0.9575367 | 0.929 | 0.598 | 2.60E-06 | 4.35E-02 |
| Cystm1 | 0.945181 | 0.929 | 0.61 | 2.89E-06 | 4.84E-02 |
| Malat1 | 0.8901726 | 1 | 0.997 | 3.29E-06 | 5.51E-02 |
| Map1b | 1.0837862 | 0.857 | 0.414 | 3.30E-06 | 5.52E-02 |
| Homer2 | 0.8413388 | 0.857 | 0.38 | 4.83E-06 | 8.09E-02 |
| Tpd52 | 0.8369763 | 1 | 0.775 | 6.81E-06 | 1.14E-01 |
| Cdo1 | 0.8450959 | 0.5 | 0.13 | 6.91E-06 | 1.16E-01 |
| Habp4 | 0.8607144 | 0.643 | 0.234 | 8.33E-06 | 1.40E-01 |
| Tubb2b | 0.8095449 | 0.857 | 0.502 | 1.07E-05 | 1.79E-01 |
| B630019K06Rik | 0.9769779 | 0.714 | 0.28 | 1.73E-05 | 2.90E-01 |
| Irf6 | 0.8546792 | 0.857 | 0.39 | 2.53E-05 | 4.24E-01 |
| Sox4 | 0.8874583 | 0.929 | 0.865 | 2.91E-05 | 4.87E-01 |
| Dcn | 1.1369661 | 0.786 | 0.35 | 2.96E-05 | 4.96E-01 |
| Sox21 | 0.9216341 | 0.857 | 0.483 | 3.36E-05 | 5.62E-01 |
| Hist3h2ba | 0.9449635 | 0.929 | 0.642 | 5.43E-05 | 9.09E-01 |
| Cd9 | 0.896098 | 1 | 0.932 | 1.17E-04 | 1.00E+00 |
| Ppp3ca | 0.7948014 | 0.857 | 0.636 | 2.02E-04 | 1.00E+00 |
| Serpinb12 | 0.8319772 | 0.643 | 0.281 | 4.89E-04 | 1.00E+00 |
| Fabp5 | 0.8698973 | 0.929 | 0.549 | 1.53E-03 | 1.00E+00 |
| Dlk1 | 1.1309571 | 0.357 | 0.141 | 9.54E-03 | 1.00E+00 |
